# Genomic insights into local scale evolution of ocular *Chlamydia trachomatis* strains within and between individuals in Gambian trachoma-endemic villages

**DOI:** 10.1101/2023.12.11.570761

**Authors:** Ehsan Ghasemian, Nkoyo Faal, Harry Pickering, Ansumana Sillah, Judith Breuer, Robin L. Bailey, David Mabey, Martin J. Holland

**Affiliations:** Department of Clinical Research, London School of Hygiene & Tropical Medicine, London, United Kingdom; Medical Research Council Unit The Gambia at London School of Hygiene and Tropical Medicine, Banjul, The Gambia; National Eye Health Programme, Ministry of Health, Kanifing, The Gambia; Division of Infection and Immunity, University College London, London, United Kingdom

**Keywords:** Trachoma, *Chlamydia trachomatis*, Whole-Genome Sequencing (WGS), Evolution

## Abstract

Trachoma, a neglected tropical disease caused by *Chlamydia trachomatis* (Ct) serovar A-C, is the leading infectious cause of blindness worldwide. Its impact is substantial, with approximately 1.9 million individuals suffering from visual impairment. Africa bears the highest burden, accounting for over 86% of global trachoma cases. We investigated Ct serovar A (SvA) and serovar B (SvB) whole genome sequences prior to the induction of mass antibiotic drug administration in The Gambia. Here, we explore the factors contributing to Ct strain diversification and implications for Ct evolution within the context of ocular infection.

A cohort study in 2002-2003 collected ocular swab samples across nine Gambian villages during a six-month follow-up study. To explore the genetic diversity of Ct within and between individuals, we conducted Whole-Genome Sequencing (WGS) on a limited number (n=43) of Ct positive samples. WGS was performed using target enrichment with SureSelect and Illumina paired-end sequencing.

Out of 43 WGS samples, 41 provided sufficient quality for further analysis. *omp*A analysis revealed that 11 samples had a highest identity to Ct strain A/HAR13 (NC_007429) and 30 had a highest identity to Ct strain B/Jali20 (NC_012686). While SvB genome sequences formed two distinct village-driven subclades, the heterogeneity of SvA sequences led to the formation of many individual branches within the Gambian SvA subclade. Comparing the Gambian SvA and SvB sequences with their reference strains; Ct A/HAR13 and Ct B/Jali20 indicated a single nucleotide polymorphism accumulation rate of 2.4 × 10^-5/site/year for the Gambian SvA and 1.3 × 10^-5/site/year for SvB variants (*p*<0.0001). Variant calling resulted in a total of 1,371 single nucleotide variants (SNVs) with a frequency > 25% in SvA sequences, and 438 SNVs in SvB sequences, from which 740 (62.8%), and 241 (66.4%) were non-synonymous, respectively. Of note, in SvA variants, highest evolutionary pressure was recorded on genes responsible for host cell modulation and intracellular survival mechanisms, whereas in SvB variants this pressure was mainly on genes essential for DNA replication/ repair mechanisms and protein synthesis. A comparison of the sequences between observed separate infection events (4-20 weeks between infections) suggested that the majority of the variations accumulated in genes responsible for host-pathogen interaction such as CTA_0166 (phospholipase D-like protein), CTA_0498 (TarP), and CTA_0948 (deubiquitinase).

This comparison of Ct SvA and SvB variants within a trachoma endemic population focused on their local evolutionary adaptation. We found a different variation accumulation pattern in the Gambian SvA chromosomal genes compared with SvB hinting at the potential of Ct serovar-specific variation in diversification and evolutionary fitness. These findings may have implications for optimizing trachoma control and prevention strategies.

**Impact Statement:** *Chlamydia trachomatis* (Ct) is a globally significant pathogen. It is the leading infectious cause of blindness—a disease called trachoma. In addition, Ct causes the majority of bacterial sexually transmitted infections. Current control measures for trachoma are based on the “SAFE” strategy; Surgery for trichiasis (S), Antibiotics (A), Facial cleanliness (F) and Environmental improvement (E). Whilst this strategy has achieved remarkable success, the target date for the global elimination of blinding trachoma as a public health problem has been pushed back from 2020 to 2030. Prior studies demonstrated evidence indicating variations in infection loads and severity among different ocular Ct serovars. However, there remains a significant knowledge gap regarding the specific genes and mechanisms responsible for these variations. We generated genetic data from two main serovars of Ct that infect human eyes: Serovar A (SvA) and Serovar B (SvB) variants collected from four villages in two different administrative regions on opposing sides of the river Gambia to elucidate (*i*) the factors driving the diversification of ocular Ct strains; (*ii*) disparities in mutation frequency/accumulation profiles; (*iii*) selective pressures between serovar A and B; and (*iv*) the dynamics of mutation accumulation within the Gambian ocular Ct positive population over a short timeframe. Our findings suggest a different variation accumulation pattern in SvA chromosomal genes compared with SvB hinting at the potential of Ct serovar-specific variation in diversification and evolutionary fitness. These findings may have implications for optimizing trachoma control and prevention strategies.

## Introduction

Trachoma, a neglected tropical disease, is the leading infectious cause of blindness worldwide, affecting marginalized populations in low-resource settings [1,2]. Trachoma is primarily caused by *Chlamydia trachomatis* (Ct) serovars A-C, with serovars A (SvA) and B (SvB) being the most commonly associated with ocular infection in Africa [3,4]. Worldwide trachoma is responsible for the visual impairment or blindness of about 1.9 million people [5–7]. Currently, an estimated 115.7 million people are at risk [6,7]. The highest concentrations of this neglected disease include 42 countries in Africa, the Middle East, Asia, and Central and South America, along with Australia. Africa has over 86% of the world’s known trachoma cases [7].

There are clear disparities in tissue tropism, disease outcome and growth rates among ocular, urogenital (UGT), and lymphogranuloma venereum (LGV) strains that are attributed to key variations in virulence genes, including *omp*A, *tar*P, *pmp*s, *trp*AB, the cytotoxin locus and *inc*A, although the genomes are highly conserved and evolutionary mechanisms have only been partially explained [8–14]. Several comparative genomics investigations have contributed novel insights into the genetic diversity and evolution of Ct [10,11,15–19]. For instance, Ct is known to undergo homologous recombination and acquire point mutations that affect tissue tropism and virulence [10,11,15]. There is cumulative evidence that implicate a family of proteins unique to *Chlamydiae*, the polymorphic membrane proteins (Pmps), in promoting niche - specific adhesion [20,21]. A study by Gomes *et al*. [22] revealed that LGV strains carry specific amino acid substitutions in PmpB, C, D and G that distinguish them from non - LGV strains, and differences in Pmp E, F and H that segregate ocular from UGT and LGV strains. In a Whole-Genome Sequencing (WGS) study focused on UGT Ct strains E and F, substantial genetic variations were identified, particularly in coding sequences related to membrane proteins such as *pmp* E and F, Type III secreted proteins (T3SS), and the cytotoxin locus, which support the assumption of higher evolutionary variability of genes involved in interactions with the host [16].

Within trachoma populations, a comparative WGS analysis of Ct strains collected from Sudanese trachoma patients [15] indicated minimal genomic diversity within this specific population. However, analyzing the genome phylogeny of the 12 Ct SvA strains from the study revealed a distinctive subclade within the larger trachoma lineage, likely stemming from an evolutionary bottleneck. Notably, three genes, namely CTA_0172, CTA_0173, and CTA_0482, exhibited extensive allelic variation, suggesting that altered expression or activity of these genes may impact the growth and survival of these ocular strains [15]. Furthermore, in a study conducted by Pickering *et al*. [3] involving trachoma patients from Amhara, Ethiopia, polymorphisms near the *omp*A locus, combined with heightened *omp*A diversity, were linked to village-level Trachomatous Inflammation-Follicular (TF) and increased Ct infection prevalence at the district level, respectively.

Prior findings by West *et al*. [23], Last *et al*. [24], and Solomon *et al*. [25] on trachoma patients suggested that the infection load might be an essential factor in the transmission of infection. Several studies on trachoma endemic communities showed that higher Ct loads were associated with Trachomatous Inflammation - Intense (TI). These studies demonstrated a link between higher Ct loads and increasing severity of inflammation in the conjunctiva [24,26,27]. Previously, a study by Ghasemian *et al*. [28] on Moroccan trachoma patients showed a significantly higher load of Ct in patients infected with SvB compared with those infected with SvA. However, there remains a significant knowledge gap regarding the specific genes and mechanisms responsible for variations in infection loads among different ocular Ct serovars. We generated genetic data from 11 SvA and 30 SvB variants collected from four villages in two administrative regions on opposing sides of the river Gambia to elucidate (*i*) the factors driving the diversification of ocular Ct strains; (*ii*) disparities in mutation frequency and accumulation profiles; (*iii*) selective pressures between SvA and SvB; and (*iv*) the dynamics of mutation accumulation within the Gambian ocular Ct positive population over a short timeframe. This study was done in the context of an investigation of Ct infection-induced immune responses and protection in trachoma [29,30], with Ct molecular diagnosis and *omp*A-sequencing the priority for the extracted DNA [31–33]. Nevertheless, an opportunistic selection of samples from this prospective cohort study allowed us to analyze Ct genomes from individuals that repeatedly tested positive over the study period, and sheds light on both immediate and long-term evolutionary trends within SvA and SvB strains from The Gambia.

## Methods

### Ethics

The samples were collected and archived under the following approvals: The joint scientific and ethics committee of the Gambian Government-Medical Research Council Gambia Unit and the London School of Hygiene & Tropical Medicine (MRC SCC: 745/781; MRC SCC L2008.75; LSHTM: 535). The study was conducted in accordance with the principles of the Declaration of Helsinki. Community leaders provided verbal consent, while written informed consent was acquired from the guardians of all study participants. In this context, a signature or thumbprint was considered an acceptable form of consent. At the time of consent, archive and secondary use were included for their potential use in pathogen genotyping studies.

### Sample collection

Sample collection was explained in detail by Barton *et al*. [29]. Briefly, for the initial screening, nine villages were chosen based on information from the Gambian National Eye Care Program (NECP), which conducted a trachoma rapid assessment survey in the Western and North Bank Regions, identifying villages where active trachoma was approximately 20% in school-age children. The study involved 345 children aged 4 to 15 from 31 family compounds, who were visited from 0 to 28 weeks every two weeks. Trachoma was graded using the WHO simplified grading systems for clinical signs by the same team of experienced trachoma graders [34]. The study occurred before mass drug administration (MDA) for trachoma control by The Gambia’s National Eye Health Programme (NEHP). Therefore, children with intense inflammatory trachoma were treated upon diagnosis, and at the study’s end, all household members were offered oral azithromycin treatment. During each visit, conjunctival swabs were collected and stored at -20°C in RNAlater.

### Detection, quantification and Whole-Genome Sequencing of *Chlamydia trachomatis*

Ct infection was assessed using a real-time polymerase chain reaction (PCR) assay targeting the 16S ribosomal RNA (rRNA), following a previously established protocol [35]. Quantification of plasmid Open Reading Frame (pORF)2 and *omc*B load was done using a droplet digital PCR (ddPCR) technique as described previously [36].

Clinical samples with an *omc*B load ≥ 10 genome equivalents were selected and WGS data were obtained directly from these samples as previously described [3]. Briefly, DNA baits spanning the length of the Ct genome were compiled by SureDesign and synthesized by SureSelectXT (Agilent Technologies, UK). DNA was sequenced at University College London/ University College London Hospitals Biomedical Research Pathogen Genomics Unit using Illumina paired-end technology (Illumina GAII or HiSeq 2000).

### Trimming, merging and quality control of the sequences

We utilized BBDuk version 38.84 to remove adapters, sequences shorter than 35 bp, and those with a Phred quality score below 20 [37]. Additionally, for merging paired reads, we employed BBMerge version 38.84 [37]. Subsequently, trimmed sequences were assessed using FastQC version 0.12.1 for various quality metrics, including “per base sequence quality”, “per sequence quality scores”, “per sequence GC content”, “per base N content”, sequence length distribution, and the presence of overrepresented sequences and adaptors [38].

### Sequence assembly and genotyping

Short reads from 43 samples were subjected to a *de novo* assembly using VELVET in conjunction with VelvetOptimiser [39]. The resulting contigs were then mapped to Ct A/HAR13 using Minimap2, specifically employing the Long-read spliced alignment data type [40,41]. This facilitated the generation of a consensus sequence, which subsequently served as a foundation for *omp*A genotyping of each sample.

### Sequence mapping and annotation

To avoid ambiguity in read mapping, we employed a masking approach for the second copy of the two largest repetitive regions of Ct reference sequences: 16S rRNA_2 and 23S rRNA_2 genes. For mapping short reads, we adopted a genovar-specific strategy, employing Bowtie2 against the masked reference genomes: Ct A/HAR13 and B/Jali20, with a minimum read identity of 90% and a minimum coverage of 10 [42]. To establish a reliable quality threshold for WGS data, we defined “good quality” as achieving a minimum coverage of 10× across at least 95% of the Ct reference genome. Chromosomal genes were defined using the annotated genome from Ct strain A/HAR13 and B/Jali20. Annotation of the consensus sequences was done in Geneious using a BLAST-like algorithm to search for best match annotations with a minimum of 80% similarity, by aligning the full length of each annotation. Antimicrobial resistance mapping using the ABRicate database was used to identify antimicrobial resistance genes in the whole genomes from Bowtie2 mapping [43,44].

### Single nucleotide polymorphism/ variant calling in individual sequences

For individual samples, “Geneious variations/SNPs caller” tool was employed to detect single nucleotide polymorphisms (SNPs) among the mapped short reads against the reference genomes: A/HAR13 or B/Jali20. The utilized parameters were as follows: a minimum coverage threshold of 10, a minimum variant frequency threshold of 80%, a maximum acceptable variant P-value of 10^-6, and a strand-bias P-value threshold of 10^-5, applied only when bias exceeded 65% (Table 1).

**Table 1.** Specific terminology to this manuscript.

| Term | Explanation |
| --- | --- |
| SNP | In this context, a Single Nucleotide Polymorphism (SNP) is characterized as a genomic position where the allele frequency is lower than 0.2 when compared to the reference chromosome of *Chlamydia trachomatis* strains A/HAR13 or B/Jali20. |
| SNV | A single nucleotide variant (SNV) was assigned to a population where the reference allele frequency was in the 10-90% range. |
| Established SNV | An established single nucleotide variant (SNV) was assigned to a population at a given site where the reference allele frequency was at least 25%. |

### Patterns of single nucleotide variant accumulation within the genes and populations

To analyze the frequency of the single nucleotide variant (SNV) within SvA compared to the SvB population, we used consensus sequences in FASTQ format to align against their respective reference strains: A/HAR13 or B/Jali20. A SNV was assigned to SvA or SvB population where the reference allele frequency was in the 10-90% range, 90% of the sequences provided coverage, and representing a maximum acceptable variant P-value of 10^-6 (Table 1). Variations with a frequency exceeding 90% were excluded to minimize those stemming from the use of different reference strains.

To discern the genes accumulating a higher number of variations and those experiencing heightened selective pressure within the SvA population compared to the SvB population, we implemented criteria for calling established SNVs. An established SNV was assigned to a population at a given site if it met the conditions of achieving a variant frequency of at least 25%, a consensus sequence coverage of at least 90%, and representing a maximum acceptable variant P-value of 10^-6 (Table 1). By imposing this 25% threshold, our aim was to exclude variations that potentially have not become established within the population or result from sequencing errors. The ratio of non-synonymous (dN) to synonymous (dS) variations were computed and assigned to each gene utilizing the Geneious.

### Phylogenetic analysis

For the global phylogenetic analyses of Ct chromosome, genome sequences from 41 isolates and 29 reference strains were aligned using progressiveMauve (Data S2 (Table S2)) [45]. A phylogenetic tree was reconstructed using RAxML (version 8.2.11) and Generalized Time Reversible (GTR) model of evolution with a γ correction for among-site rate variation with four rate categories and 1000 bootstraps [46]. Moreover, plasmids from 41 isolates and 27 reference strains were used to build a phylogenetic tree following the same methodology (Data S2 (Table S2)). Here, for the first time, we employed the accurate Ct strain B/Tunis864 genome in drawing the Ct global phylogenetic tree. In the supplementary materials, we have included a concise explanation to address the ongoing confusion pertaining to the labeling of the whole genome sequences of Ct strains B/HAR36 and B/Tunis864 (Data S1).

Gene alignments were generated using MAFFT (version v7.490) with a 200 PAM/K = 2 scoring matrix [47,48]. PhyML was utilized to estimate maximum likelihood phylogenies of aligned sequences with a GTR model of evolution and 1000 bootstraps [49].

### Statistics

Microsoft Excel (version 16.78) and GraphPad Prism (version 10.0.3) were used for designing the graphs. A *p*-value of < 0.05 was considered to reflect a statistically significant difference. A non-parametric, two-tailed, Mann-Whitney test was performed to examine any association between Ct infection load and Ct genovar. We used a parametric, unpaired, two-tailed t-test to explore the significance of difference in the distribution of the variations in SvA than in SvB sequences.

## Results

### Sample collection, *Chlamydia trachomatis* infection and sequencing quality data

From the entire cohort collection 1019 samples tested positive for Ct at least once. After selecting and using these samples for confirmatory Ct diagnostic tests, *omp*A, toxin and *tar*P amplicon sequencing tests [30], a subset of 43 representative samples were selected that had sufficient DNA yield, quality and Ct load for WGS. After quality assessments of the whole genome sequences, 41/43 had passed defined quality control measures and achieved a minimum coverage of 10× across at least 95% of the Ct reference genome. The *omp*A genotyping process was carried out on these 41 sequences using NCBI BLAST-n, which resulted in 11 sequences exhibiting highest similarity to strain A/HAR13 (NC_007429) and 30 sequences to strain B/Jali20 (NC_012686). These 41 samples were derived from 26 participants and originated from four villages (Fig. 1). Among these individuals, 15 sampling time-points (A-N) were documented, and the study samples were ultimately composed of three samples from one patient, two samples from 13 patients, and one sample from 12 patients (Table 2). The selected samples originated from participants with a mean age of 8.8 years, consisting of 17 males and 9 females, with 22 presenting clinical signs of TF, while four displayed no clinical signs meeting the WHO simplified grading score definition of trachoma (Table 2 and Data S2 (Table S1)).

**Fig. 1.**
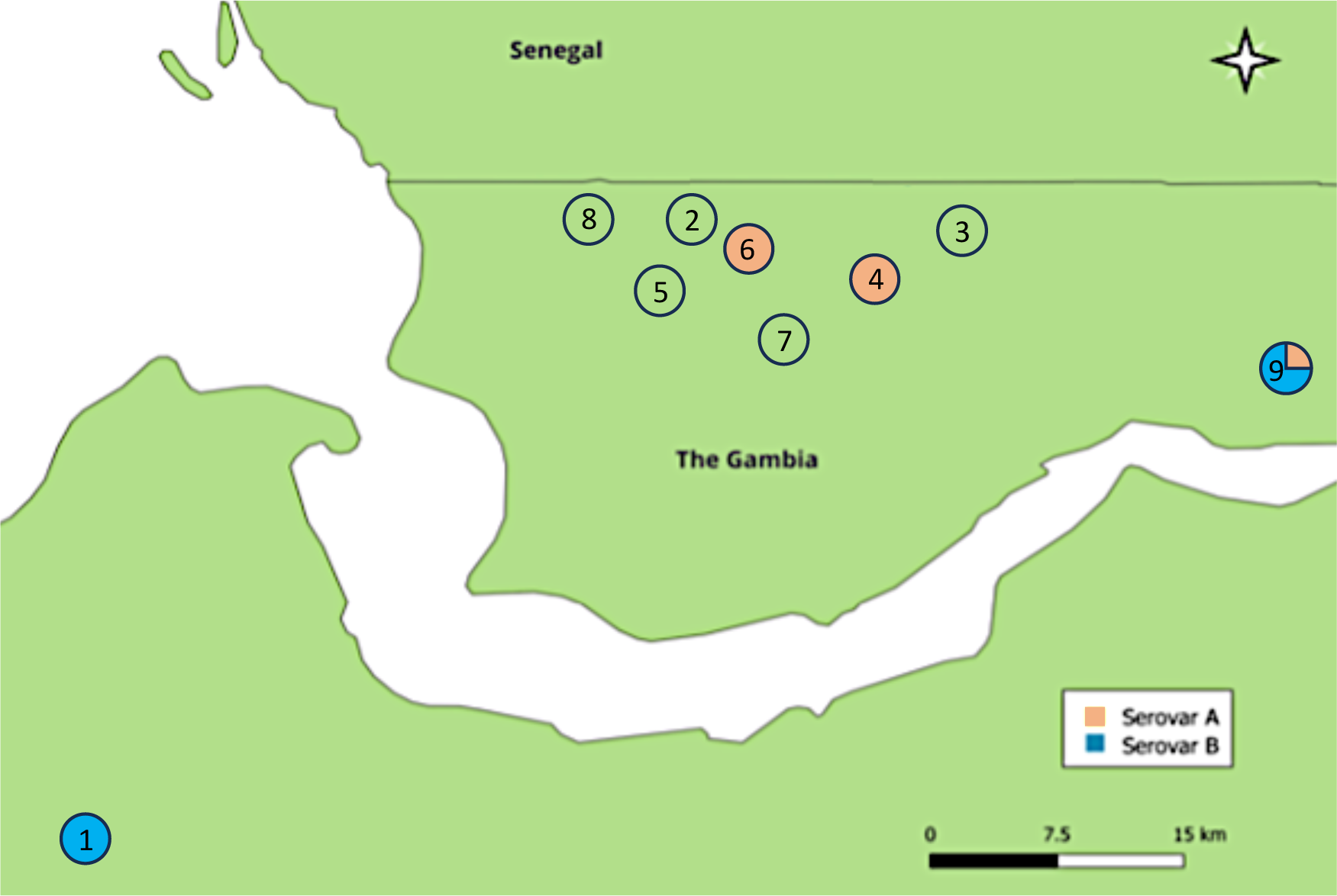
Sampling sites and geographical distribution of *Chlamydia trachomatis omp*A gene among the Gambian samples. In total nine villages in The Gambia were included in the sampling process from which 41 samples from villages 1, 4, 6 and 9 provided good quality whole-genome sequences that were included in this study. Red color indicates the presence of *C. trachomatis* (Ct) serovar A in the village, and blue color indicates the presence of Ct serovar B in the village. Empty circles depict villages that were included in the sampling process but did not provide Ct whole genomes.

**Table 2.** Baseline demographics, trachoma grades and CT *omp*A type and *omc*B load of study participants.

| Village | Patient ID | Sex | Age | Trachoma grade | Number of infections | Time between specimen | *ompA* genotype | Plasmid genotype | *omcB* copies/μL |
| --- | --- | --- | --- | --- | --- | --- | --- | --- | --- |
| Village 1 |  |  |  |  |  |  |  |  |  |
|  | 010202D & J | F | 5 | TF | 2 | 12 weeks | B | B | 14.8 |
|  | 010306G & M | F | 5 | TF | 2 | 12 weeks<sup>s</sup> | B | B | 1628 |
|  | 010404L | F | 10 | TF | 1 | - | B | B | 37.3 |
|  | 010702D & K | F | 7 | TF | 2 | 14 weeks | B | B | 56.4 |
|  | 010703D & J | F | 4 | TF | 2 | 12 weeks | B | B | 1112.6 |
|  | 010705I | M | 10 | Normal | 1 | - | B | B | 68.8 |
|  | 011402D & I | F | 10 | TF | 2 | 10 weeks | B | B | 1960.7 |
|  | 011501C & J | F | 5 | TF | 2 | 14 weeks | B | B | 1551.2 |
|  | 011502C & J | F | 4 | TF | 2 | 14 weeks | B | B | 193.1 |
| Village 4 |  |  |  |  |  |  |  |  |  |
|  | 040104A | M | 6 | TF | 1 | - | A | A | 249.3 |
|  | 040119A & G & L | M | 13 | TF | 3 | 12 and 8 weeks | A | A | 27.1 |
| Village 6 |  |  |  |  |  |  |  |  |  |
|  | 060121A & K | M | 8 | TF & Normal | 8 | 20 weeks | A | A | 358.6 |
|  | 060123G | M | 7 | TF | 7 | - | A | A | 216 |
|  | 060124L | F | 8 | TF | 8 | - | A | A | 24.7 |
| Village 9 |  |  |  |  |  |  |  |  |  |
|  | 090109J & N | M | 15 | TF & Normal | 2 | 8 weeks | B | B | 408 |
|  | 090120N | M | 9 | Normal | 1 | - | B | B | 69.3 |
|  | 090132N | M | 13 | TF | 1 | - | A | A | 29.5 |
|  | 090134J&L | M | 9 | TF | 2 | 4 weeks | B | B | 114.3 |
|  | 090139L | M | 9 | TF | 1 | - | B | B | 34.5 |
|  | 090142N | M | 13 | TF | 1 | - | B | A | 27 |
|  | 090154J & L | M | 11 | Normal | 1 | - | B | B | 69 |
|  | 090155N | M | 9 | TF | 2 | 4 weeks | B | B | 297.3 |
|  | 090167N | M | 9 | Normal | 1 | - | B | B | 28.8 |
|  | 090168L & N | M | 10 | TF & Normal | 2 | 4 weeks | B | B | 278.2 |
|  | 090172J & M | M | 11 | TF | 2 | 6 weeks | A | A | 933.6 |
|  | 090195N | M | 8 | TF | 1 | - | B | B | 37.5 |

\* To calculate the average *omcB* gene load for samples with multiple infection time-points, we aggregated data from all time-points.
§ Among those that we obtained WGS data in more than one sampling time-point, only 010306 remained constantly PCR-positive between two sampling time-points (12 weeks).

On average, each sample generated 3,540,540 raw reads, with 639,183 merged paired-reads successfully mapped to the Ct reference genome. Across these 41 samples, the average coverage of the Ct reference genome was 113 (merged paired-reads), with a corresponding mean confidence score of 38 (Data S2 (Table S1)).

### *omp*A diversity and *omc*B copy number

Maximum BLAST-n homology (https://www.ncbi.nlm.nih.gov/blast/) against the Ct *omp*A gene assigned 11 samples to Ct SvA, strain A/HAR13 and 30 samples to Ct SvB, strain B/Jali20. Serovar distribution across villages was as follows: 16 samples were classified as SvB in village one, four samples as SvA in village four, four samples as SvA in village six, and three samples as SvA, with 15 samples as SvB in village nine (Table 2). The average *omc*B gene copy number for SvB variants (420.4 copies/μL) was higher than that for SvA variants (270.5 copies/μL), however, the difference was not significant (p=0.6597).

In line with BLAST-n results, phylogenetic analysis of *omp*A assigned the Gambian sequences into two distinct clusters (Fig. 2) where SvA sequences grouped closely with Ct strain A/HAR13 (isolation year: 1958) [50], and two SvA Gambian reference strains; A/D213 (isolation year: 2001) [51], and A/D230 (isolation year: 2001) [18]. Moreover, all three SvB reference strains from the Gambia; B/Jali16 (isolation year: 1985) [50], B/Jali20 (isolation year: 1985) [52], and B/M48 (isolation year: 2007) [18], grouped together with SvB sequences from village nine. Among sequences classified as SvB, a unique substitution was observed in sequences from village one at position 893 (C>T=A>V) of *omp*A that differentiate these sequences from SvB sequences from village nine and the Gambian SvB strains deposited previously in the ENA; B/Jali16, B/Jali20, and B/M48 [18,50,52]. Moreover, two substitutions at positions 186 (G>A=M>I) and 268 (G>A=A>T) of *omp*A were specific to the SvB sequences that originated from the Gambia including those from our study.

**Fig. 2.**
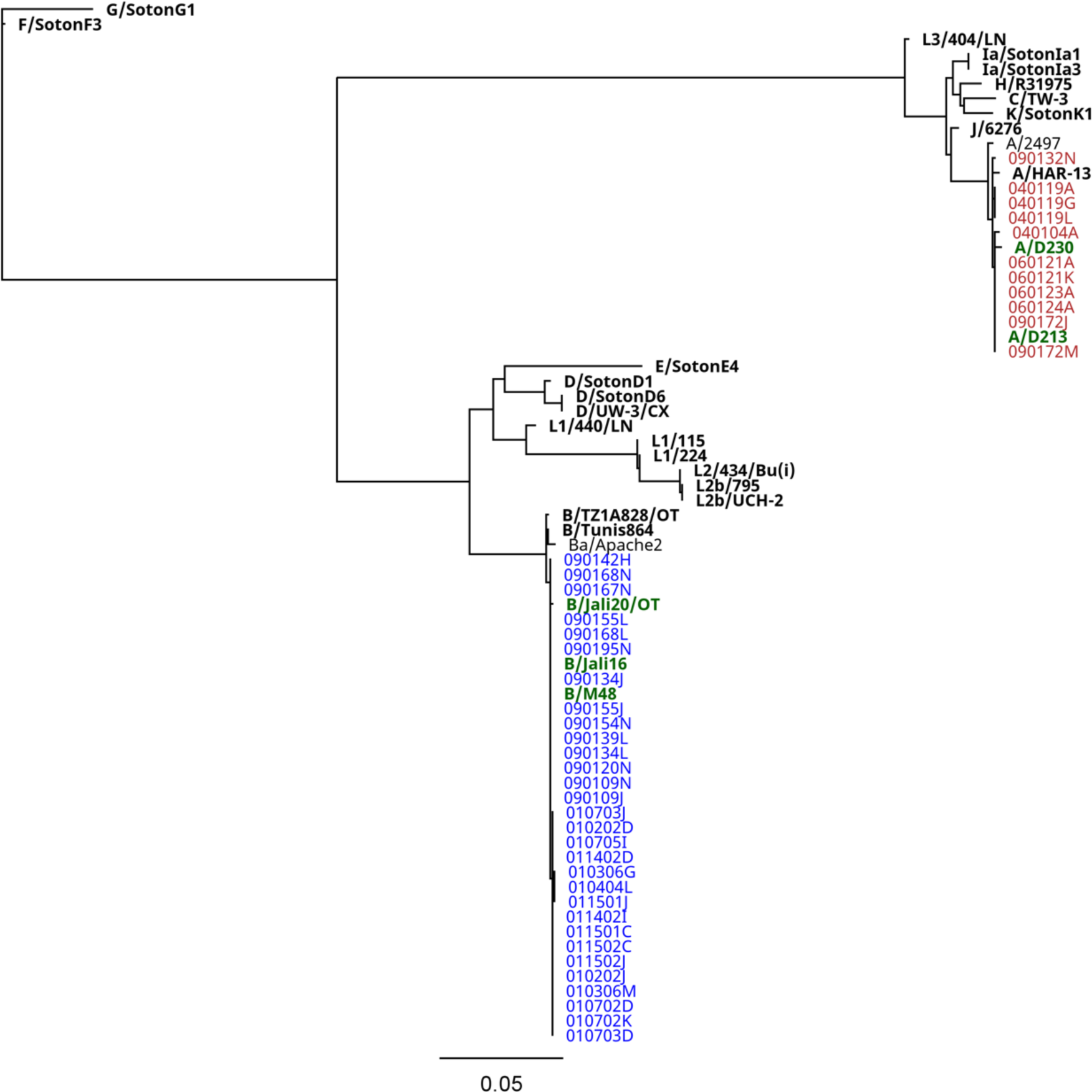
Maximum likelihood phylogenetic reconstruction of ocular *Chlamydia trachomatis omp*A sequences from the Gambia. The phylogenetic analysis encompasses 41 *C. trachomatis* (Ct) positive samples collected in The Gambia and 29 Ct reference strains. The initial two digits of each sample identifier signify the respective recruitment village. Samples testing positive for Ct serovar A are denoted in red, while those for serovar B are indicated in blue. All reference strains are highlighted in bold, with strains originating from The Gambia represented in green.

### Global phylogeny of the Gambian *Chlamydia trachomatis* genomes and plasmids

We studied the phylogenetic distribution of SvA and SvB genomes derived from our study and a selection of chlamydial genomes that corresponded to the four major Ct clades found globally, including LGV, UGT, prevalent UGT, and ocular clades [18,51]. To better resolve the phylogeny of the Gambian Ct genomes, we introduced three less familiar genome sequences from previous Gambian studies: A/D213 [35], A/D230[18], and B/M48[18], alongside the known SvB reference strains B/Jali16 [34] and B/Jali20 [36]. All Gambian genomes clustered within the ocular clade, forming two distinct subclades of SvA and SvB (Fig. 3). This suggests that the Gambian genomes were derived from at least two separate ancestral sources. Remarkably, the SvA subclade may represent a locally endemic clone, distinct from the more common SvA reference strains including A/HAR13 and A/2497 (Fig. 3).

**Fig. 3.**
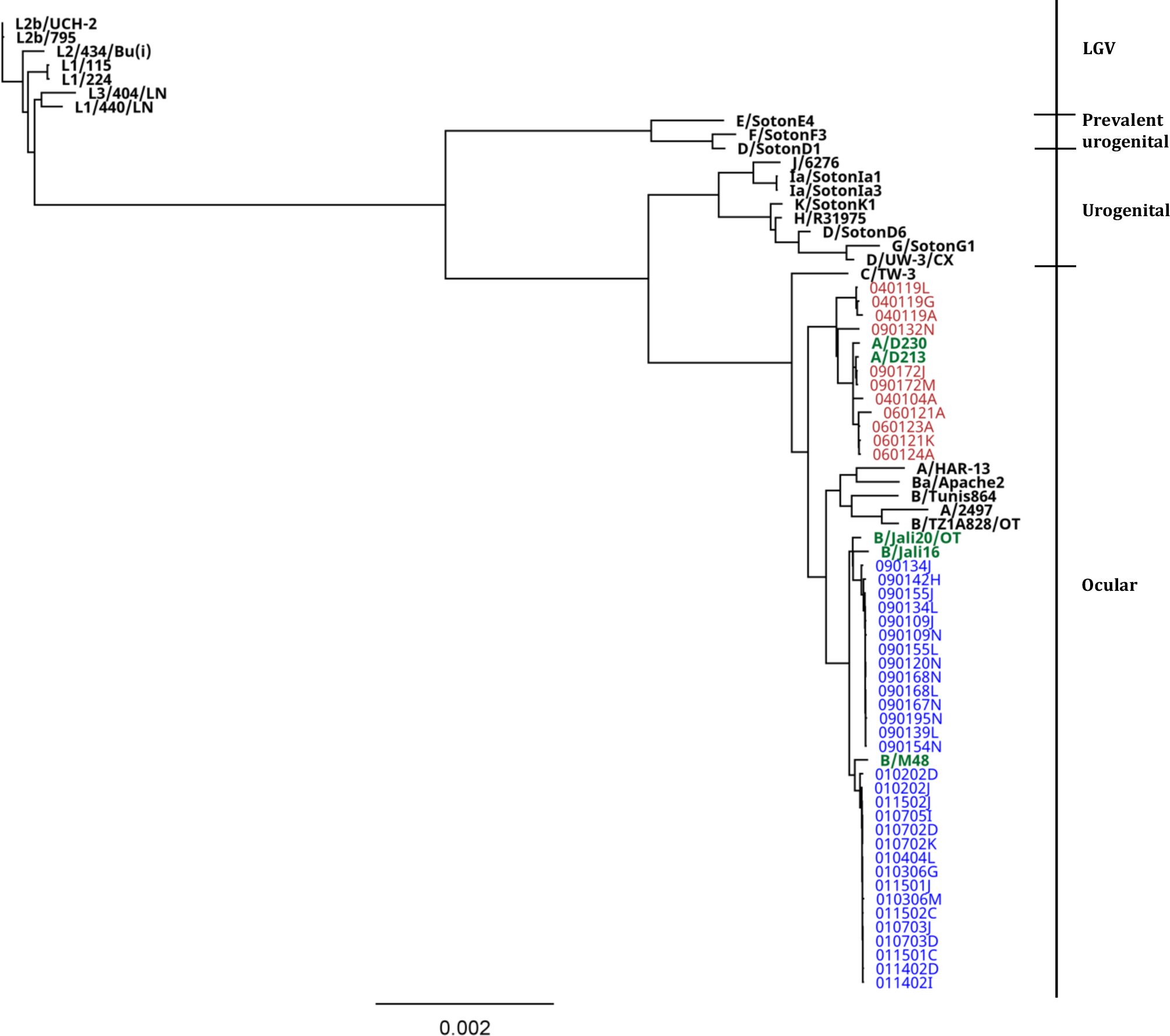
Global phylogeny of *Chlamydia trachomatis* chromosomal sequences from the Gambia. The four major *C. trachomatis* (Ct) lineages are listed on the right. Chromosomal sequences were aligned using progressiveMauve, and a phylogenetic tree was reconstructed employing RAxML, incorporating the Generalized Time Reversible (GTR) model of evolution with a γ correction for among-site rate variation, employing four rate categories, and subjected to 1000 bootstrap resampling iterations. Whole genome sequences of samples testing positive for Ct serovar A are denoted in red, while those for serovar B are indicated in blue. All reference strains are highlighted in bold, with strains originating from The Gambia represented in green.

Within the Gambian SvA subclade, we observed two distinct groups. The first comprises three genomes isolated from a single individual in village four (Fig. 3). The second group includes one genome from village four, along with genomes from village six and village nine (Fig. 3). These genomes share a common ancestor with reference strains A/D213 and A/D230. Sequences from village six form a separate subgroup within the second group (Fig. 3). Surveying the SvB subclade, we noted that sequences from village one and village nine divide into two distinct groups (Fig. 3). SvB genomes from village nine cluster together and share a common ancestor with strain B/Jali20, and SvB genomes from village one share a common ancestor with strains B/M48 and B/Jali16 (Fig. 3).

Like the Ct chromosome phylogeny, the global phylogeny of Ct plasmid reveals four distinct clusters: “LGV”, “UGT”, “prevalent UGT”, and “ocular” clades, with the exception of strain Ba/Apache2 that grouped within the “prevalent UGT” cluster (Fig. 4) [53]. Gambian plasmids formed one distinct subclade within the ocular clade from all other ocular reference strains. We identified six branches within the Gambian subclade including (*i*) sequences from Ct SvA isolates; (*ii*) sequences from SvB isolates from village nine; (*iii*) sequences from SvB isolates from village one that are sharing ancestor with the reference strain B/M48; (*iv*) reference strain B/Jali16; (*v*) reference strain B/Jali20; and (*vi*) one plasmid from village nine (090142H) that did not group with any other plasmids from village nine and formed a separate branch (Fig. 4). Intriguingly, when examining the BLAST-n homology of 090142H plasmid, it demonstrated the highest homology to plasmid of Ct strain A/2497. Looking at the Gambian Ct SvA plasmid sequences, three plasmids from one participant in village four (040119A, G, L) grouped together with one plasmid from village nine, while all other plasmids formed a separate subgroup sharing common ancestors with the reference strains A/D213 and A/D230 (Fig. 4).

**Fig. 4.**
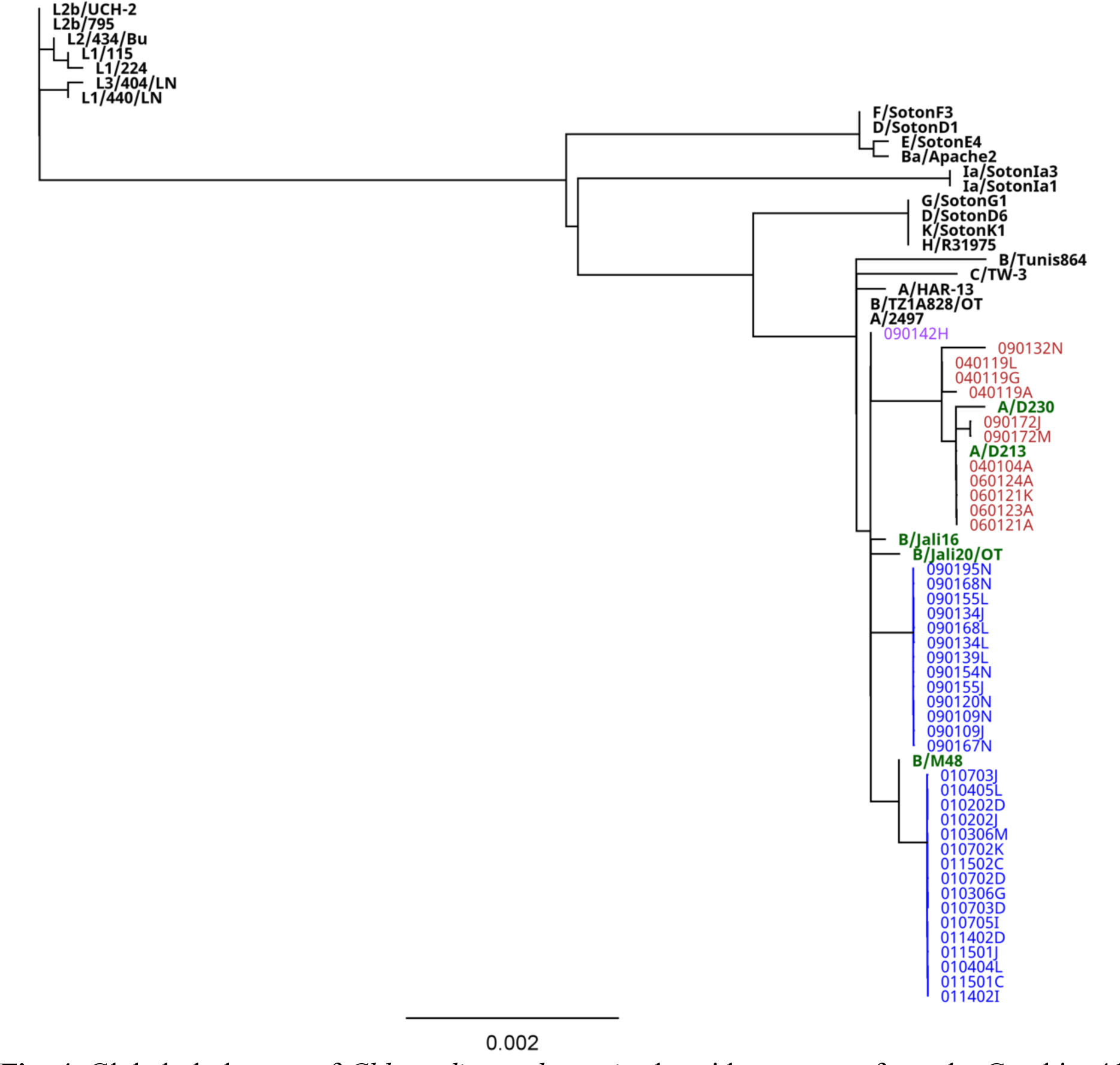
Global phylogeny of *Chlamydia trachomatis* plasmid sequences from the Gambia. 41 Gambian and 27 *C. trachomatis* (Ct) reference strains plasmid sequences were aligned using progressiveMauve, and a phylogenetic tree was reconstructed employing RAxML, incorporating the Generalized Time Reversible (GTR) model of evolution with a γ correction for among-site rate variation, employing four rate categories, and subjected to 1000 bootstrap resampling iterations. Plasmid sequences from samples testing positive for Ct serovar A are denoted in red, while those for serovar B are indicated in blue. Plasmid sequence of sample 090142H formed a separate branch from the rest of the Gambian sequences and is marked in violet. All reference strains are highlighted in bold, with strains originating from The Gambia represented in green.

Among Ct SvA plasmid sequences we found seven SNVs with a frequency of at least 25% from which three were nonsynonymous. Three (43%) out of seven variations accumulated in CDS3 (replicative DNA helicase). While four other variations accumulated in CDS1 (14.3%), CDS2 (14.3%), CDS7 (14.3%) and CDS8 (14.3%). Two out of three nonsynonymous variations accumulated in CDS3 and one in CDS1 (pgp7). In comparison, we found eight established SNVs among Ct SvB plasmid sequences from which two were nonsynonymous. The distribution of the established SNVs on the plasmid of Ct SvB isolates was as follows: two (25%) in CDS1, two (25%) in CDS2, one (12.5) in CDS3, two (25%) in CDS4 and one (12.5%) in CDS8. Nonsynonymous variations were found in CDS1 and CDS4.

### Single nucleotide polymorphism accumulation in individual sequences

We identified fixed mutations in individual samples as those where 80% of the sequencing reads with a minimum coverage of 10 disagreed with the reference base. In the case of SvA sequences, the range of SNP was between 1,117 and 1,175, while for SvB sequences, it ranged from 144 to 295 (*p*<0.0001). On average, among SvA sequences from villages four, six, and nine, there were 1,156, 1,132, and 1,169 SNPs, respectively. Conversely, among SvB sequences from villages one and nine, the average number of SNPs detected was 285 and 238, respectively (*p*<0.0001). Considering the respective years in which the samples were collected; Ct strain A/HAR13; 1958, and Ct strain B/Jali20; 1985, the SNP accumulation rates for our Gambian SvA and SvB variants were calculated as ∼2.5 × 10^-5 and ∼1.4 × 10^-5 /site/year, respectively (*p*<0.0001) [50,52].

### Mutation frequency among the Gambian variants

To assess the frequency of the variations within the sequences classified as SvA and SvB, a variation was assigned where the reference allele frequency was in the 10-90% range. Based on the variation frequency, we categorized them into three groups: (*i*) those with a frequency between 10-25% were considered not established at the village level; (*ii*) SNVs with a frequency between 25-60% were deemed established in one village; and (*iii*) SNVs with a frequency between 60-90% were regarded as established in at least two villages. For both SvA and SvB populations the majority of the SNVs (52.1% and 92.9%, respectively) were established in one village (Fig. 5). For SvA population while 34.3% of the SNVs were in common among the residents of at least two villages (Fig. 5a and 5b), only 2.7% of the SNVs in SvB population could get established in both village one and nine (Fig. 5c and 5d). Furthermore, we noted a higher number of unestablished SNVs at a village level in the SvA (13.5%) population compared to the SvB (4.4%) population (Fig. 5).

**Fig. 5.**
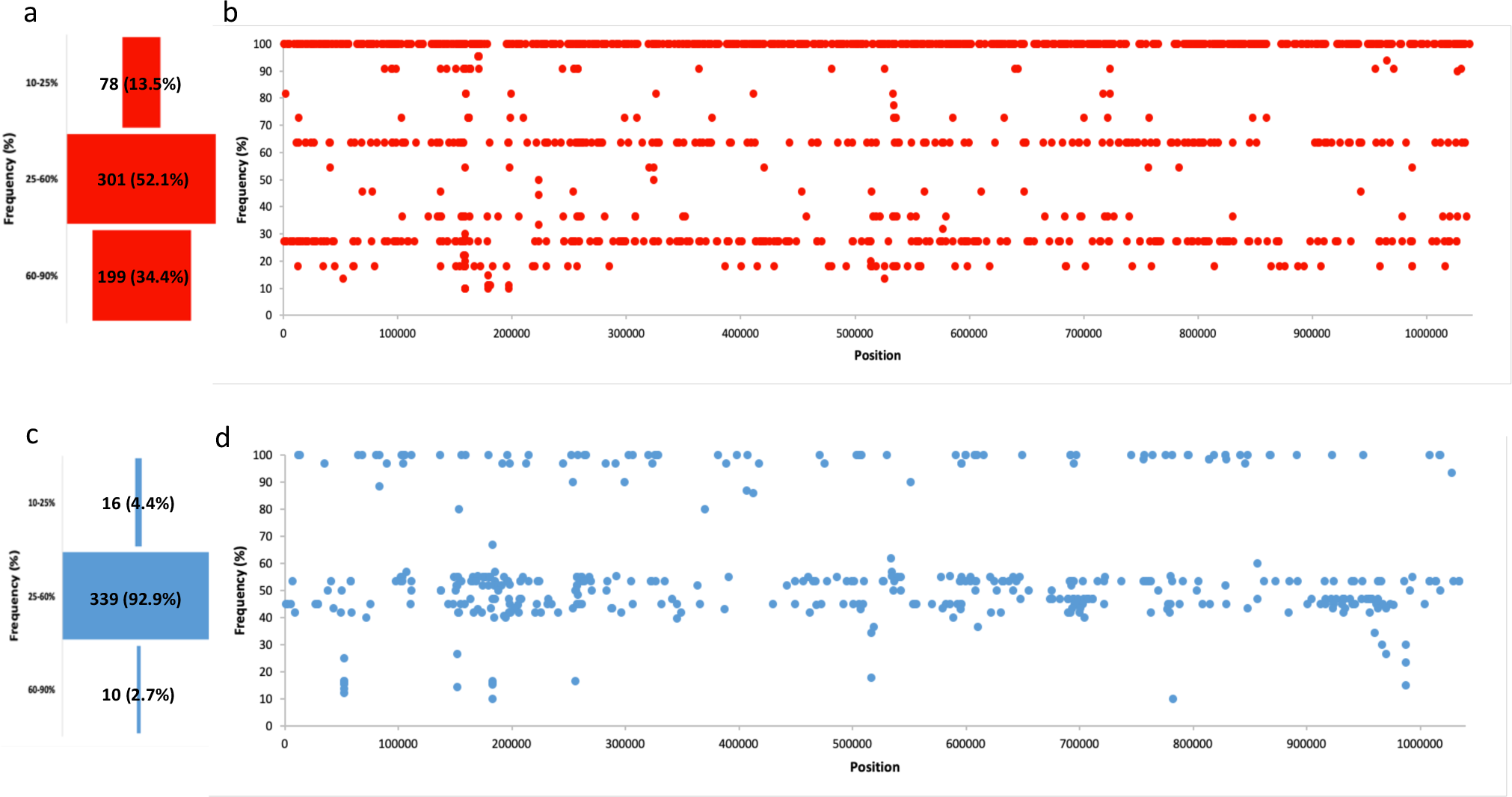
Variation frequency analysis in the Gambian *Chlamydia trachomatis* variants. In panels (a) and (b), the red funnel and dot plot graphs display the frequency of single nucleotide variant (SNV) accumulated in the Gambian serovar A strains in relation to the reference strain A/HAR13. Conversely, in panels (c) and (d), the blue funnel and dot plot graphs illustrate the frequency of SNVs accumulated in the Gambian serovar B strains in comparison to the reference strain B/Jali20. The funnel exclusively incorporates SNVs with a frequency ranging from 10% to 90%.

### Gambian serovar B strains appear under higher selective pressure than serovar A strains

Analysing variations with a frequency of at least 25% yielded a total of 1,371 established SNVs among SvA sequences and 438 established SNVs among SvB sequences. Among the SvA sequences, 740 established SNVs (62.8%) were categorized as nonsynonymous. In contrast, within the SvB sequences, 241 (66.4%) were designated as nonsynonymous. This observation suggests a potentially higher selective pressure acting on SvB variants compared to SvA variants within the Gambia.

### *Chlamydia trachomatis* genes under highest selective pressure

While the established SNV rates on the chromosomes of SvA and SvB variants are ∼13.2 and ∼4.2/10 kbp, respectively, it is noteworthy that the plasticity zone (PZ), spanning from *acc*B to *trp*A, exhibits a heightened established SNV rate for both SvA (∼20.2 mutations/10 kbp) and SvB (∼10.4 mutations/10 kbp). The SvA PZ dN/dS ratio (∼1.7) was like other regions of the SvA chromosome (∼1.7) whereas the SvB PZ dN/dS ratio (∼4) was higher than the SvB chromosome (∼2) and more than double that of SvA. This may be indicative of a higher selective pressure on the PZ of Gambian SvB variants than SvA variants. Furthermore, we recorded 5 established SNVs (4 nonsynonymous and 1 synonymous) in *omp*A of SvA sequences and 2 established SNVs (2 nonsynonymous) in *omp*A of SvB sequences. A variation at position 268 (G>A=A>T) of the Gambian SvB *omp*A located on variable domain one. Examining the genes that have accumulated the highest number of established SNVs and dN/dS ratio in Ct SvA sequences (Fig. 6a, Fig. 6b, Data S2 (Table S3, Table S4)), we can categorize these genes based on their functional significance as follows: (*i*) genes involved in energy/nutrition/protein transport and trafficking pathways including CTA_0070/*npt*1, CTA_0087 (T3SS), CTA_0146, CTA_0156, CTA_0241 (YitT), CTA_0251, CTA_0390/*bio*Y, CTA_RS02415/*sec*D, and CTA_0747/*suf*D; (*ii*) genes pivotal for Ct virulence, such as CTA_0310/*inc*A, CTA_0498/*tar*P, CTA_0675, CTA_0948 (deubiquitinase (DUB)); (*iii*) genes important for Ct membrane structure, exemplified by CTA_0062, CTA_0271, and CTA_0389; and (*iv*) those important for Ct DNA replication and repair, represented by CTA_0155/*lig*A [50,54–65].

**Fig. 6.**
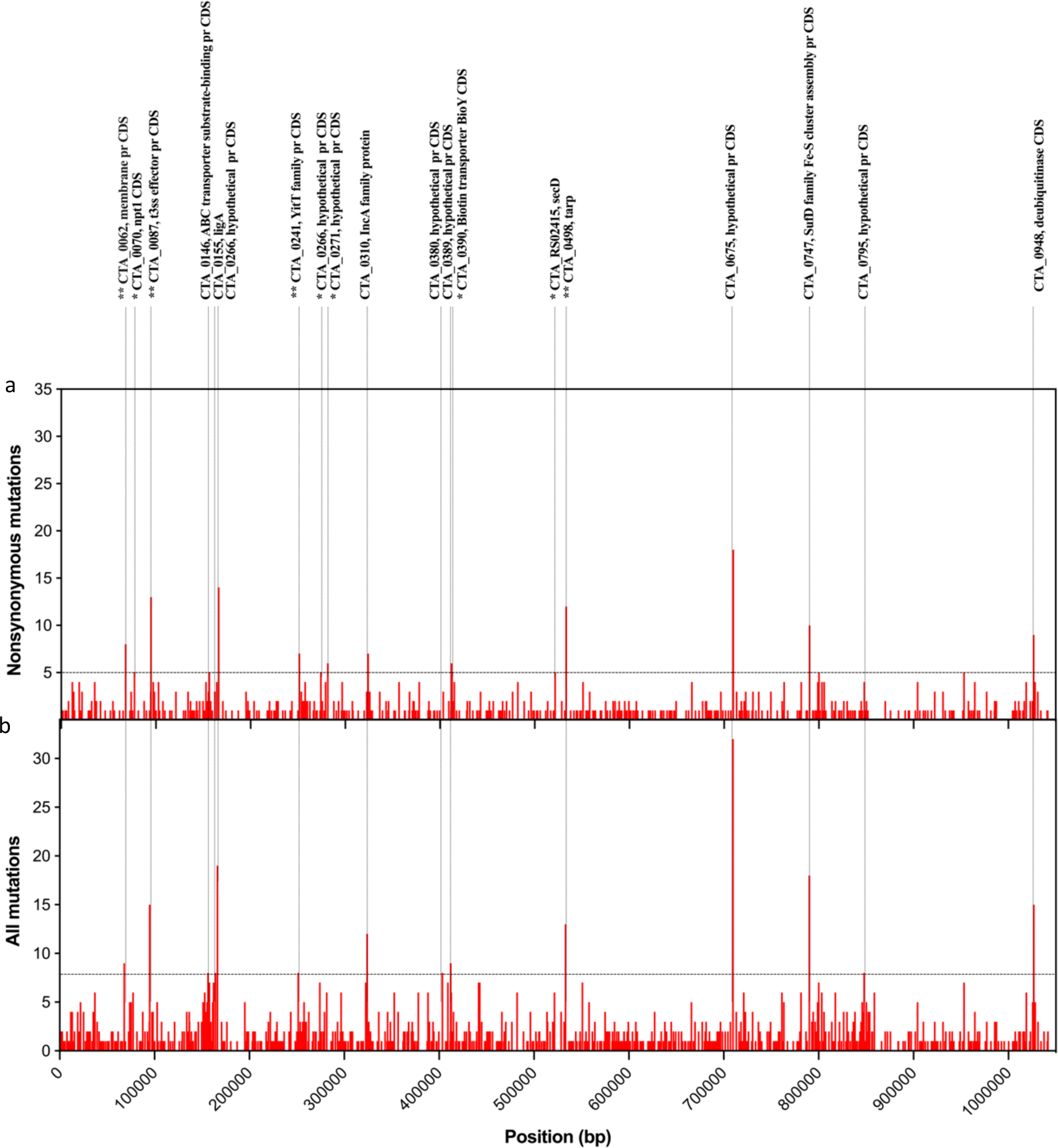
Gene-Level variation accumulation profiles in *Chlamydia trachomatis* SvA variants. (a) The graph shows the total number of variant sites (upper bars) detected for each *C. trachomatis* gene, and (b) the number of nonsynonymous substitution sites (lower bars) exceeding a frequency 25% among the Gambian SvA sequences. The chromosomal genes are ordered according to genome annotation of the A/HAR13 strain (NC_007429). The genes showing at least five nonsynonymous substitution sites (and no synonymous variant sites) or a ratio of nonsynonymous (dN)/synonymous (dS) equal or above five (Top 1%) are labeled above the graph. The genes showing at least eight total established single nucleotide variant (SNV) sites (Top 1%) are labeled above the graph. Single asterisk (*) labels genes with the highest ratio of dN/dS (Top 1%). Double asterisk (**) labels genes that were in common between those with the highest ratio of dN/dS, and highest number of established SNV sites (Top 1%).

SvB genes with the highest established SNVs counts and dN/dS ratios are distributed among four categories (Fig. 7a, Fig. 7b, Data S2 (Table S3, Table S4)). The largest group encompasses (*i*) genes involved in DNA replication, repair, and RNA synthesis, notably JALI_0951/*inf*B, JALI_5251/*rps*C, JALI_6121 (UvrD helicase), JALI_6131, JALI_7991/*dna*G [66–70]. The other groups include (*ii*) genes relevant to amino acid synthesis and metabolic processes, such as JALI_1641/*trp*B, JALI_1771, JALI_6361, and JALI_8241/*glm*S; (*iii*) genes associated with Ct virulence, including JALI_RS01230/*inc*A, JALI_4581/*tar*P, and JALI_8191/*pmp*D; and (*iv*) genes participating in transport and trafficking pathways, exemplified by JALI_2291 [57,60,71–76].

**Fig. 7.**
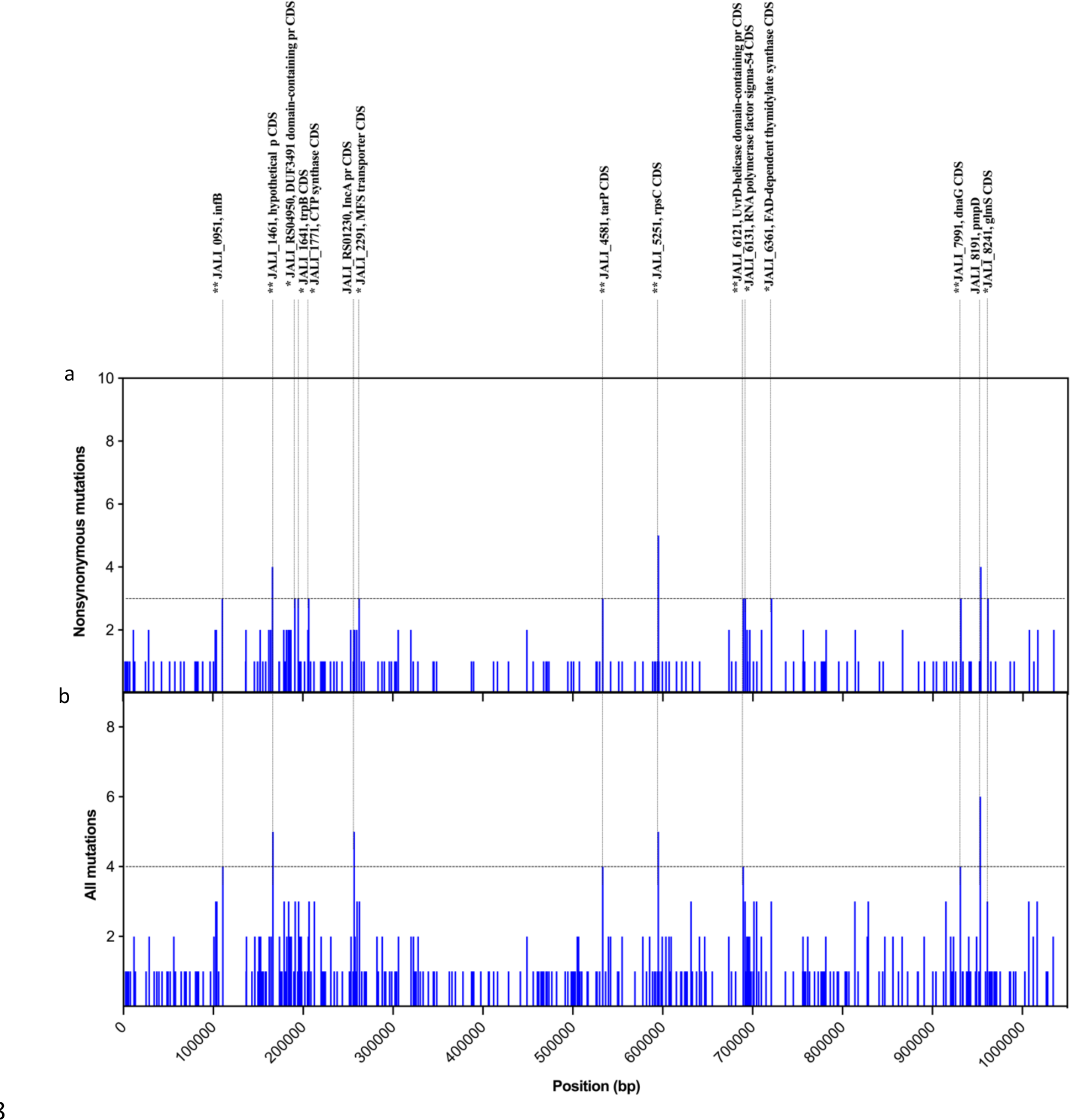
Gene-Level variation accumulation profiles in *Chlamydia trachomatis* SvB variants. (a) The graph shows the total number of variant sites (upper bars) detected for each *C. trachomatis* gene, and (b) the number of nonsynonymous substitution sites (lower bars) exceeding a frequency 25% among the Gambian SvB sequences. The chromosomal genes are ordered according to genome annotation of the B/Jali20 strain (NC_012686). (a) The genes showing at least three nonsynonymous substitution sites (and no synonymous variant sites) or a ratio of nonsynonymous (dN)/synonymous (dS) equal or above three (Top 1%) are labeled above the graph. (b) The genes showing at least four total established single nucleotide variant (SNV) sites (Top 1%) are labeled above the graph. Single asterisk (*) labels genes with the highest ratio of dN/dS (Top 1%). Double asterisk (**) labels genes that were in common between those with the highest ratio of dN/dS, and highest number of SNV sites (Top 1%).

### Tryptophan operon analyses revealed a truncation in TrpB

Maximum likelihood phylogenetic analysis of the *trp*AB genes resulted in the formation of a separate clade specific to the ocular strains (Data S2 (Fig. S1)). Within this ocular clade, the Gambian *trp*AB sequences can be further divided into three distinct subclades: (*i*) comprises SvB sequences only from village one, along with the reference strain B/M48; (*ii*) encompasses seven SvA sequences derived from villages four, six, and nine, in addition to the reference strains A/D213 and A/D230; and (*iii*) consists of SvB sequences from village nine, which cluster together with four SvA sequences from villages four and nine (Data S2 (Fig. S1)).

Examination of the *trp*B gene has revealed the presence of an insertion spanning positions 1,315 to 1,316. This insertion event resulted in a frameshift, leading to the early termination of the *trp*B CDS. Notably, this mutation is conserved across all SvB sequences originating from village one and is also present in the reference strain B/M48 (Data S2 (Fig. S2)).

### Short-term mutation accumulation trends in *Chlamydia trachomatis* SvA and SvB

We obtained WGS data from three individuals who tested repeatedly positive for SvA in villages four, six, and nine, as well as 11 individuals who tested repeatedly positive for SvB in villages one and nine. On average, the time interval between the first and second identified infections used for WGS for SvA sequences was ∼12.7 weeks, while for SvB sequences, it was ∼9.8 weeks. A comparison between the first and second infection within each participant revealed a total of 171 SNPs events (33 among SvA sequences and 138 among SvB sequences), resulting in an average of 11 SNPs/SvA genome and 11.5 SNPs/SvB genome. Notably, we identified 35 SNPs within Ct PZ. This represents an elevated SNP rate, averaging at 5.8 SNPs/10 kb, which is nearly four times higher than the average observed rate (1.6 SNPs/10 kb) across the entire Ct genome.

Data S2 (Table S5) and Fig. 8 represent top 1% of Ct genes with the highest number of mutation events collected between second and first infections. Sequence comparisons between the second and first infection revealed four categories of the genes that have accumulated majority of the SNPs including (*i*) genes associated with Ct virulence such as CTA_0498/*tar*P, CTA_0166 (phospholipase D-like protein (PDL)), CTA_0948 (DUB); (*ii*) CTA_0021/*ile*S associated with amino acid metabolic process; (*iii*) CTA_0484/*omc*A associated with Ct extracellular matrix; and (*iv*) CTA_0140 that is involved in transport and trafficking pathways (Fig. 8 and Data S2 (Table S5)) [57,64,77–80].

**Fig. 8.**
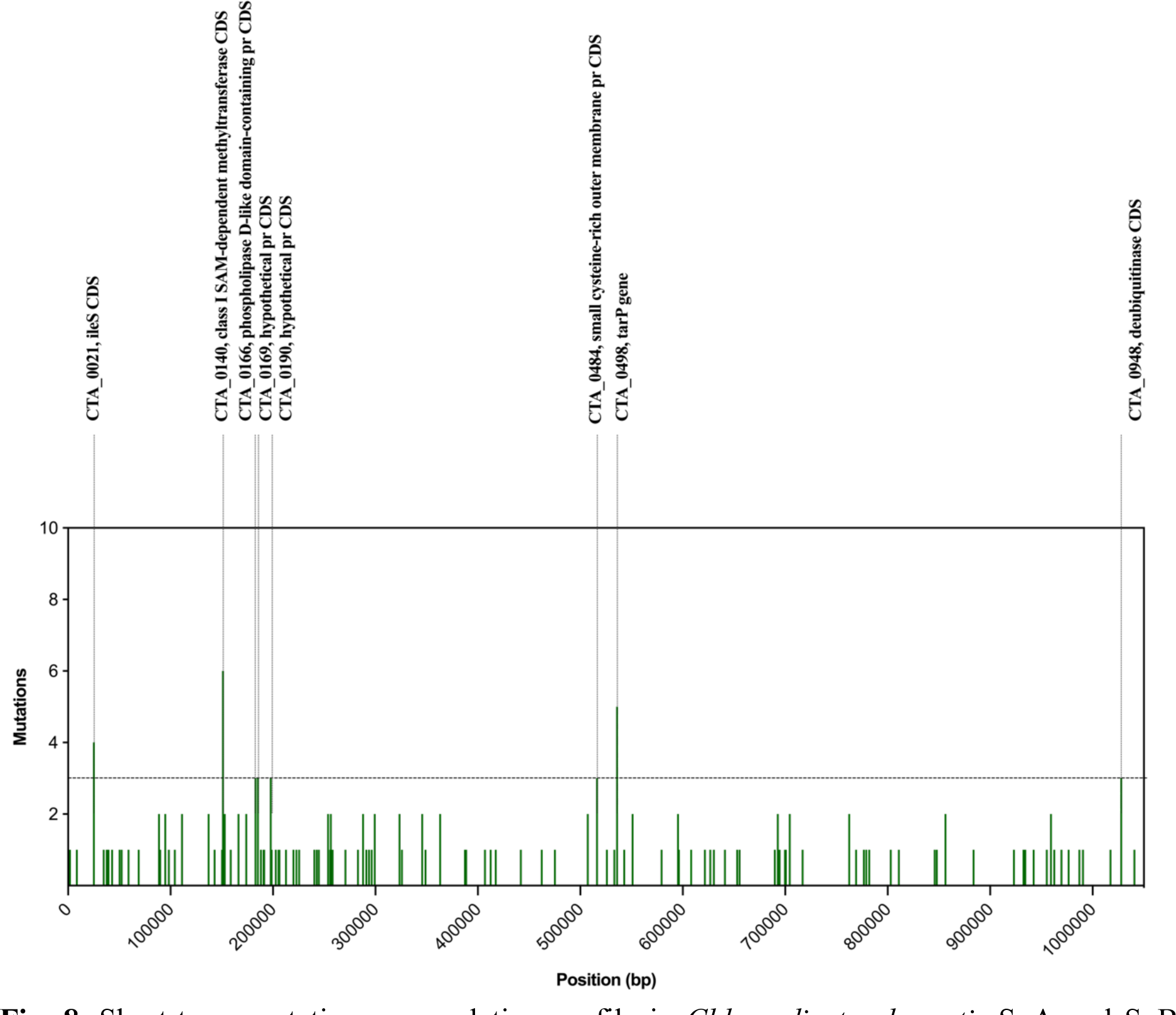
Short-term mutation accumulation profile in *Chlamydia trachomatis* SvA and SvB sequences. Each column represents the cumulative mutation count within each *C. trachomatis* gene across three participants who tested positive for serovar A and 11 participants who tested positive for serovar B in the second compared to the first infection timepoint. The analysis average timespan was 11.25 weeks. The chromosomal genes are ordered according to genome annotation of the A/HAR13 strain (NC_007429).

## Discussion

We conducted WGS on Ct isolates obtained from 11 SvA and 30 SvB variants from The Gambia. Of note, samples positive for Ct SvA originated from three villages (village 4 (Jokadu), 6 (Lower Niumi), and 9 (Central Baddibu)) in the North Bank Region of the river and samples positive for Ct SvB originated from two villages, one in the North Bank Region (village 9) and one in the West Coast Region (village 1 (Kombo South)). Through phylogenetic analysis of Ct *omp*A gene sequences, we showed that diversification in the Gambian SvA *omp*A sequences is not driven by the location of the villages, whereas for SvB sequences there is a distinction between sequences from village one and nine. Of note, both SvA and SvB o*mp*A sequences grouped closely with Ct A/HAR13 and B/Jali20, sequences collected ∼50 and ∼20 years prior to this study [50,52]. Previously, studies on Ct UGT strains showed that the expansion of genotype E, currently the most prevalent UGT genotype, may be due to increased fitness at or around *omp*A, preventing recombination being fixed in this region, being simply a stochastic increase, or being a combination of the two [18].

Of note, differences in the frequency of the SNVs among SvA and SvB sequences support the global phylogeny of the Gambian Ct chromosomes. There is a higher influx of the unestablished SNVs (ranging from 10% to 25%) among SvA population (13.5%) than among SvB population (4.4%). This may explain the higher phylogenetic diversity observed within the Gambian SvA sequences in contrast to the SvB sequences. Our findings on SvB sequences support a prior observation by Alkhidir *et al.* [15] suggesting a similar phylogenetic relatedness for two SvB Gambian isolates collected ∼20 years apart: B/Jali20 and B/M48, indicating slow and geographically related diversification. The majority of the SNVs for both SvA (52.1%) and SvB (92.9%) sequences were established on a village level (ranging from 25% to 60%). This prevalence of village-level SNVs among SvB sequences likely contributes to the formation of distinct clusters within SvB sequences, corresponding to their respective villages of origin. This supports previous findings in trachoma endemic populations that suggest geographical clustering of ocular Ct strains [13,18,81]. Moreover, while 34.4% of the SNVs could reach a frequency of 60-90% and become established among the majority of SvA population, potentially accounting for the emergence of a distinctive Gambian SvA subclade within the traditional ocular Ct clade, only 2.7% of the SNVs among SvB population could reach to a frequency of 60-90%.

In line with prior studies that demonstrated concordance of chromosome and plasmid phylogeny [11,51,82,83], the phylogenetic position of the Gambian plasmids, with the exception of one sample, is consistent with the whole genome phylogeny. BLAST-n analysis of sample 09142H plasmid showed the highest homology to Ct SvA strain A/2497, while *omp*A and chromosome phylogeny classified this sample as SvB. Previous studies on UGT strains presented rare evidence of horizontal plasmid transfer events, recombination events and plasmid swapping [18,82,84]. While our data reveals a strong association between the chromosomal genotype and plasmid that suggests their co-evolution, there remains a possibility of recombination or a swapping event for the plasmid of 090142H.

Prior data for Ct LGV strains and *Chlamydia psittaci* estimated substitution rates of 2.1 × 10^-7 and 1.7 × 10^-4 SNPs/site/year [18,85,86]. Our results estimated 2.5 × 10^-5 and 1.4 × 10^- 5 SNPs/site/year for the Gambian SvA and SvB variants, respectively [50,52]. This suggests an almost two times higher SNP accumulation rate among the Gambian SvA sequences compared with SvB sequences, and approximately 100-200x higher substitution rate in the Gambian ocular strains than that reported for Ct LGV [18,85]. This higher rate can be in part due to the differences in Ct serovars and might also be driven by acquisition and recombination events between Ct strains in the Gambian populations [18,85]. The dN/dS ratio is a measure of selective pressure [87], higher ratios indicate positive selection [88,89]. We found a slightly higher percentage of nonsynonymous substitutions in SvB (66.4%, dN/dS=2) compared with SvA (62.8% dN/dS=1.7) sequences. Previously, Borges and Gomes [90] and Seth-Smith *et al*. [85] reported 55% and 75% nonsynonymous substitutions in protein coding regions of Ct LGV strains, respectively.

In agreement with prior data [28], we observed a higher Ct load for SvB compared with SvA variants. Previously, Kari *et al*. [91] provided evidence demonstrating a direct relationship between polymorphisms in specific Ct genes and virulence properties of trachoma strains. In SvA sequences, genes linked to host cell modulation (e.g., “DUB”), host-pathogen interactions (e.g., “T3SS effector protein”), and intracellular survival and nutrient acquisition (e.g., “*npt*1”), and in SvB sequences, genes related to RNA translation (e.g., “*inf*B”), DNA replication, (e.g., “*dna*G”), and amino acid synthesis (e.g., “*trp*B”) accumulated the highest number of established SNVs and/or ratio of dN/dS [54,55,64,65]. These genes in SvB strains potentially lead to increased replicative fitness of Ct within the host cell [66,70,92]. A study conducted by Sigalova *et al*. [93] classified chlamydial genes into three functional groups based on annotated Clusters of Orthologous Genes. Combining this classification with our observations, indicates that a larger proportion of “core genes” among SvB than SvA variants experience greater evolutionary pressure. Conversely, a higher number of “periphery genes” among SvA compared to SvB variants are subject to increased selective pressure. We reported an insertion in the *trp*B gene of SvB sequences from village one, that causes a frameshift in *trp*B, and therefore the truncation of TrpB. These results support prior findings indicating high evolutionary pressure on *trp* operons of Ct [19,94].

Our study is subject to several limitations that warrant consideration. Limited participants with repeated SvA infections challenge understanding of short-term SNP accumulation patterns in SvA variants. A larger sample size from wider geographic regions in the Gambia would enhance the robustness of our findings. In addition, it is worth acknowledging that the choice of A/HAR13 as a reference strain collected in 1958 in Egypt for SvA sequences and B/Jali20 as a reference strain collected in 1985 in the Gambia for SvB sequences may have introduced some bias into the variation frequency and accumulation data among SvA and SvB populations [50,52].

In conclusion, while Gambian SvB variants appear to be well-adapted to the population and need little further adaptation in order to maintain themselves within the Gambian population, SvA variants exhibit a propensity for diversification, accumulating new mutations. Our findings suggest that geographical factors play a role in driving the diversification and adaptation of Ct strains, highlighting the impact of geospatial differences on Ct evolution. Previously, differences in the rate of mutation accumulation in the chromosome of Ct strains was explained by the influence of various factors, including adaptation dynamics, different mutation accumulation and repair speed, geographical specifications, and the influence of mass community-level treatment [15,24,27,51,95–98]. We emphasize that a more extensive investigation in a larger trachoma-endemic population, involving participants over an extended timeframe, is necessary to investigate our speculation suggesting different mutation accumulation rates in the chromosome of ocular Ct strains. Finally, our observations imply that the degree of evolutionary pressure on ocular Ct strains may vary, reflecting the specific fitness of each Ct strain, as manifested in the specific genes experiencing the highest evolutionary pressure in the Gambian SvA compared to SvB variants.

## Ethical approval and consent to participate

The samples were collected and archived under the following approvals: The joint scientific and ethics committee of the Gambian Government-Medical Research Council Gambia Unit and the London School of Hygiene & Tropical Medicine (MRC SCC: 745/781; MRC SCC L2008.75; LSHTM: 535). The study was conducted in accordance with the principles of the Declaration of Helsinki. Community leaders provided verbal consent, while written informed consent was acquired from the guardians of all study participants.

## Acknowledgments

We would like to thank the study participants, as well as the trachoma field and technical teams, including Omar Manneh, Omar Camara, Hassan Joof, Pateh Makalo, Esther Aryee, Isatou Sarr, Mass Laye. Special thanks to Dr Julius Schachter and Jeanne Moncada for providing us with the original stocks of Ct strains B/HAR36 and B/Tunis864.

## Author contribution

Conceived study: H.P. M.J.H. A.S. DCWM, RLB. Conducted fieldwork: N.F., A.S., M.J.H., R.L.B. Conducted laboratory analyses: N.F., H.P, MJH. Conducted data analysis and interpretation: E.G., H.P., NF and MJH. Drafted manuscript: E.G., M.J.H. Commented, edited, approved manuscript: all.

## Data Summary

Gambian *Chlamydia trachomatis* sequencing data in the form of fastq.gz files used in this study can be accessed from the European Nucleotide Archive (ENA) project accession PRJEB68379 (Accession ERR12330790-ERR12330830). Reference strains B/Tunis864 and B/HAR36 sequencing data and assemblies in the form of fastq.gz and bam files can be accessed from the European Nucleotide Archive (ENA) project accession PRJEB68374 (Accession ERR12253485-ERS16770979). All packages used for data analysis are linked to citations.

## References

1. Solomon, A.W.; Burton, M.J.; Gower, E.W.; Harding-Esch, E.M.; Oldenburg, C.E.; Taylor, H.R.; Traoré, L. Trachoma. Nat Rev Dis Primers 2022, 8, 32, doi:10.1038/s41572-022-00359-5.

2. Taylor, H.R.; Burton, M.J.; Haddad, D.; West, S.; Wright, H. Trachoma. Lancet 2014, 384, 2142–2152, doi:10.1016/S0140-6736(13)62182-0.

3. Pickering, H.; Chernet, A.; Sata, E.; Zerihun, M.; Williams, C.A.; Breuer, J.; Nute, A.W.; Haile, M.; Zeru, T.; Tadesse, Z.;, et al. Genomics of Ocular *Chlamydia Trachomatis* After 5 Years of SAFE Interventions for Trachoma in Amhara, Ethiopia. J Infect Dis 2022, 225, 994–1004, doi:10.1093/infdis/jiaa615.

4. Andreasen, A.A.; Burton, M.J.; Holland, M.J.; Polley, S.; Faal, N.; Mabey, D.C.; Bailey, R.L. Chlamydia Trachomatis OmpA Variants in Trachoma: What Do They Tell Us? PLoS Negl Trop Dis 2008, 2, e306, doi:10.1371/journal.pntd.0000306.

5. Wolle, M.A.; West, S.K. Ocular Chlamydia Trachomatis Infection: Elimination with Mass Drug Administration. Expert Rev Anti Infect Ther 2019, 17, 189–200, doi:10.1080/14787210.2019.1577136.

6. Trachoma; WHO, 2022. Available Online: https://www.who.int/News-Room/Fact-Sheets/Detail/Trachoma (Accessed on 05 October 2022).

7. WHO Alliance for the Global Elimination of Trachoma: Progress Report on Elimination of Trachoma, 2022. Wkly Epidemiol Rec 2023, 28, 297–314.

8. Abdelsamed, H.; Peters, J.; Byrne, G.I. Genetic Variation in Chlamydia Trachomatis and Their Hosts: Impact on Disease Severity and Tissue Tropism. Future Microbiol 2013, 8, 1129–1146, doi:10.2217/fmb.13.80.

9. Hadfield, J.; Bénard, A.; Domman, D.; Thomson, N. The Hidden Genomics of Chlamydia Trachomatis. Curr Top Microbiol Immunol 2018, 412, 107–131, doi:10.1007/82_2017_39.

10. Joseph, S.J.; Didelot, X.; Gandhi, K.; Dean, D.; Read, T.D. Interplay of Recombination and Selection in the Genomes of Chlamydia Trachomatis. Biol Direct 2011, 6, 28, doi:10.1186/1745-6150-6-28.

11. Joseph, S.J.; Didelot, X.; Rothschild, J.; de Vries, H.J.C.; Morré, S.A.; Read, T.D.; Dean, D. Population Genomics of Chlamydia Trachomatis: Insights on Drift, Selection, Recombination, and Population Structure. Mol Biol Evol 2012, 29, 3933–3946, doi:10.1093/molbev/mss198.

12. Somboonna, N.; Mead, S.; Liu, J.; Dean, D. Discovering and Differentiating New and Emerging Clonal Populations of Chlamydia Trachomatis with a Novel Shotgun Cell Culture Harvest Assay. Emerg Infect Dis 2008, 14, 445–453, doi:10.3201/eid1403.071071.

13. Last, A.R.; Pickering, H.; Roberts, C. h.; Coll, F.; Phelan, J.; Burr, S.E.; Cassama, E.; Nabicassa, M.; Seth-Smith, H.M.B.; Hadfield, J.;, et al. Population-Based Analysis of Ocular Chlamydia Trachomatis in Trachoma-Endemic West African Communities Identifies Genomic Markers of Disease Severity. Genome Med 2018, 10, 15, doi:10.1186/s13073-018-0521-x.

14. Carlson, J.H.; Hughes, S.; Hogan, D.; Cieplak, G.; Sturdevant, D.E.; McClarty, G.; Caldwell, H.D.; Belland, R.J. Polymorphisms in the Chlamydia Trachomatis Cytotoxin Locus Associated with Ocular and Genital Isolates. Infect Immun 2004, 72, 7063–7072, doi:10.1128/IAI.72.12.7063-7072.2004.

15. Alkhidir, A.A.I.; Holland, M.J.; Elhag, W.I.; Williams, C.A.; Breuer, J.; Elemam, A.E.; El Hussain, K.M.K.; Ournasseir, M.E.H.; Pickering, H. Whole-Genome Sequencing of Ocular Chlamydia Trachomatis Isolates from Gadarif State, Sudan. Parasit Vectors 2019, 12, 518, doi:10.1186/s13071-019-3770-7.

16. Eder, T.; Kobus, S.; Stallmann, S.; Stepanow, S.; Köhrer, K.; Hegemann, J.H.; Rattei, T. Genome Sequencing of Chlamydia Trachomatis Serovars E and F Reveals Substantial Genetic Variation. Pathog Dis 2017, 75, doi:10.1093/femspd/ftx120.

17. Beder, T.; Saluz, H.P. Virulence-Related Comparative Transcriptomics of Infectious and Non-Infectious Chlamydial Particles. BMC Genomics 2018, 19, doi:10.1186/s12864-018-4961-x.

18. Hadfield, J.; Harris, S.R.; Seth-Smith, H.M.B.; Parmar, S.; Andersson, P.; Giffard, P.M.; Schachter, J.; Moncada, J.; Ellison, L.; Vaulet, M.L.G.;, et al. Comprehensive Global Genome Dynamics of Chlamydia Trachomatis Show Ancient Diversification Followed by Contemporary Mixing and Recent Lineage Expansion. Genome Res 2017, 27, 1220– 1229, doi:10.1101/gr.212647.116.

19. Somboonna, N.; Ziklo, N.; Ferrin, T.E.; Suh, J.H.; Dean, D. Clinical Persistence of Chlamydia Trachomatis Sexually Transmitted Strains Involves Novel Mutations in the Functional Αββα Tetramer of the Tryptophan Synthase Operon. mBio 2019, 10, doi:10.1128/mBio.01464-19.

20. Elwell, C.; Mirrashidi, K.; Engel, J. Chlamydia Cell Biology and Pathogenesis. Nat Rev Microbiol 2016, 14, 385–400, doi:10.1038/nrmicro.2016.30.

21. Nunes, A.; Gomes, J.P. Evolution, Phylogeny, and Molecular Epidemiology of Chlamydia. Infection, Genetics and Evolution 2014, 23, 49–64, doi:10.1016/j.meegid.2014.01.029.

22. Almeida, F.; Borges, V.; Ferreira, R.; Borrego, M.J.; Gomes, J.P.; Mota, L.J. Polymorphisms in Inc Proteins and Differential Expression of Inc Genes among Chlamydia Trachomatis Strains Correlate with Invasiveness and Tropism of Lymphogranuloma Venereum Isolates. J Bacteriol 2012, 194, 6574–6585, doi:10.1128/JB.01428-12.

23. West, E.S.; Munoz, B.; Mkocha, H.; Holland, M.J.; Aguirre, A.; Solomon, A.W.; Bailey, R.; Foster, A.; Mabey, D.; West, S.K. Mass Treatment and the Effect on the Load of Chlamydia Trachomatis Infection in a Trachoma-Hyperendemic Community. Invest Ophthalmol Vis Sci 2005, 46, 83–87, doi:10.1167/iovs.04-0327.

24. Last, A.; Burr, S.; Alexander, N.; Harding-Esch, E.; Roberts, C.H.; Nabicassa, M.; Cassama, E.; Mabey, D.; Holland, M.; Bailey, R. Spatial Clustering of High Load Ocular Chlamydia Trachomatis Infection in Trachoma: A Cross-Sectional Population-Based Study. Pathog Dis 2017, 75, doi:10.1093/femspd/ftx050.

25. Solomon, A.W.; Holland, M.J.; Alexander, N.D.E.; Massae, P.A.; Aguirre, A.; Natividad-Sancho, A.; Molina, S.; Safari, S.; Shao, J.F.; Courtright, P.;, et al. Mass Treatment with Single-Dose Azithromycin for Trachoma. N Engl J Med 2004, 351, 1962–1971, doi:10.1056/NEJMoa040979.

26. Nash, S.D.; Chernet, A.; Moncada, J.; Stewart, A.E.P.; Astale, T.; Sata, E.; Zerihun, M.; Gessese, D.; Melak, B.; Ayenew, G.;, et al. Ocular Chlamydia Trachomatis Infection and Infectious Load among Pre-School Aged Children within Trachoma Hyperendemic Districts Receiving the SAFE Strategy, Amhara Region, Ethiopia. PLoS Negl Trop Dis 2020, 14, e0008226, doi:10.1371/journal.pntd.0008226.

27. Harding-Esch, E.M.; Holland, M.J.; Schémann, J.-F.; Sillah, A.; Sarr, B.; Christerson, L.; Pickering, H.; Molina-Gonzalez, S.; Sarr, I.; Andreasen, A.A.;, et al. Impact of a Single Round of Mass Drug Administration with Azithromycin on Active Trachoma and Ocular Chlamydia Trachomatis Prevalence and Circulating Strains in The Gambia and Senegal. Parasit Vectors 2019, 12, 497, doi:10.1186/s13071-019-3743-x.

28. Ghasemian, E.; Inic-Kanada, A.; Collingro, A.; Mejdoubi, L.; Alchalabi, H.; Keše, D.; Elshafie, B.E.; Hammou, J.; Barisani-Asenbauer, T. Comparison of Genovars and Chlamydia Trachomatis Infection Loads in Ocular Samples from Children in Two Distinct Cohorts in Sudan and Morocco. PLoS Negl Trop Dis 2021, 15, e0009655, doi:10.1371/journal.pntd.0009655.

29. Barton, A.; Rosenkrands, I.; Pickering, H.; Faal, N.; Harte, A.; Joof, H.; Makalo, P.; Ragonnet, M.; Olsen, A.W.; Bailey, R.L.;, et al. A Systems Serology Approach to the Investigation of Infection-Induced Antibody Responses and Protection in Trachoma. Front Immunol 2023, 14, doi:10.3389/fimmu.2023.1178741.

30. Lutter, E.I.; Bonner, C.; Holland, M.J.; Suchland, R.J.; Stamm, W.E.; Jewett, T.J.; McClarty, G.; Hackstadt, T. Phylogenetic Analysis of Chlamydia Trachomatis Tarp and Correlation with Clinical Phenotype. Infect Immun 2010, 78, 3678–3688, doi:10.1128/IAI.00515-10.

31. Barton, A.; Faal, N.; Ramadhani, A.; Derrick, T.; Mafuru, E.; Mtuy, T.; Massae, P.; Malissa, A.; Joof, H.; Makalo, P.;, et al. Longitudinal Changes in Tear Cytokines and Antimicrobial Proteins in Trachomatous Disease. PLoS Negl Trop Dis 2023, 17, e0011689, doi:10.1371/journal.pntd.0011689.

32. Faal, N.; Bailey, R.L.; Jeffries, D.; Joof, H.; Sarr, I.; Laye, M.; Mabey, D.C.W.; Holland, M.J. Conjunctival FOXP3 Expression in Trachoma: Do Regulatory T Cells Have a Role in Human Ocular Chlamydia Trachomatis Infection? PLoS Med 2006, 3, e266, doi:10.1371/journal.pmed.0030266.

33. Holland, M.J.; Faal, N.; Sarr, I.; Joof, H.; Laye, M.; Cameron, E.; Pemberton-Pigott, F.; Dockrell, H.M.; Bailey, R.L.; Mabey, D.C.W. The Frequency of Chlamydia Trachomatis Major Outer Membrane Protein-Specific CD8+ T Lymphocytes in Active Trachoma Is Associated with Current Ocular Infection. Infect Immun 2006, 74, 1565– 1572, doi:10.1128/IAI.74.3.1565-1572.2006.

34. Thylefors, B.; Dawson, C.R.; Jones, B.R.; West, S.K.; Taylor, H.R. A Simple System for the Assessment of Trachoma and Its Complications. Bull World Health Organ 1987, 65, 477–483.

35. Burton, M.J.; Holland, M.J.; Jeffries, D.; Mabey, D.C.W.; Bailey, R.L. Conjunctival Chlamydial 16S Ribosomal RNA Expression in Trachoma: Is Chlamydial Metabolic Activity Required for Disease to Develop? Clin Infect Dis 2006, 42, 463–470, doi:10.1086/499814.

36. Roberts, C.H.; Last, A.; Molina-Gonzalez, S.; Cassama, E.; Butcher, R.; Nabicassa, M.; McCarthy, E.; Burr, S.E.; Mabey, D.C.; Bailey, R.L.;, et al. Development and Evaluation of a Next-Generation Digital PCR Diagnostic Assay for Ocular Chlamydia Trachomatis Infections. J Clin Microbiol 2013, 51, 2195–2203, doi:10.1128/JCM.00622-13.

37. Bushnell, B.; Rood, J.; Singer, E. BBMerge – Accurate Paired Shotgun Read Merging via Overlap. PLoS One 2017, 12, e0185056, doi:10.1371/journal.pone.0185056.

38. Andrews, S. FastQC: A Quality Control Tool for High Throughput Sequence Data. 2010, Available Online: http://www.bioinformatics.babraham.ac.uk/Projects/Fastqc/.

39. Zerbino, D.R.; Birney, E. Velvet: Algorithms for de Novo Short Read Assembly Using de Bruijn Graphs. Genome Res 2008, 18, 821–829, doi:10.1101/gr.074492.107.

40. Li, H. New Strategies to Improve Minimap2 Alignment Accuracy. Bioinformatics 2021, 37, 4572–4574, doi:10.1093/bioinformatics/btab705.

41. Li, H. Minimap2: Pairwise Alignment for Nucleotide Sequences. Bioinformatics 2018, 34, 3094–3100, doi:10.1093/bioinformatics/bty191.

42. Langmead, B.; Salzberg, S.L. Fast Gapped-Read Alignment with Bowtie 2. Nat Methods 2012, 9, 357–359, doi:10.1038/nmeth.1923.

43. Seemann, T. ABRicate: Mass Screening of Contigs for Antiobiotic Resistance Genes. 2016, Available Online: https://github.com/Tseemann/Abricate.

44. Zankari, E.; Hasman, H.; Cosentino, S.; Vestergaard, M.; Rasmussen, S.; Lund, O.; Aarestrup, F.M.; Larsen, M. V. Identification of Acquired Antimicrobial Resistance Genes. Journal of Antimicrobial Chemotherapy 2012, 67, 2640–2644, doi:10.1093/jac/dks261.

45. Darling, A.C.E.; Mau, B.; Blattner, F.R.; Perna, N.T. Mauve: Multiple Alignment of Conserved Genomic Sequence with Rearrangements. Genome Res 2004, 14, 1394– 1403, doi:10.1101/gr.2289704.

46. Stamatakis, A. RAxML Version 8: A Tool for Phylogenetic Analysis and Post-Analysis of Large Phylogenies. Bioinformatics 2014, 30, 1312–1313, doi:10.1093/bioinformatics/btu033.

47. Katoh, K. MAFFT: A Novel Method for Rapid Multiple Sequence Alignment Based on Fast Fourier Transform. Nucleic Acids Res 2002, 30, 3059–3066, doi:10.1093/nar/gkf436.

48. Katoh, K.; Standley, D.M. MAFFT Multiple Sequence Alignment Software Version 7: Improvements in Performance and Usability. Mol Biol Evol 2013, 30, 772–780, doi:10.1093/molbev/mst010.

49. Guindon, S.; Dufayard, J.-F.; Lefort, V.; Anisimova, M.; Horik, W.; Gascuel, O. New Algorithms and Methods to Estimate Maximum-Likelihood Phylogenies: Assessing the Performance of PhyML 3.0. Syst Biol 2010, 59, 307–321, doi:10.1093/sysbio/syq010.

50. Carlson, J.H.; Porcella, S.F.; McClarty, G.; Caldwell, H.D. Comparative Genomic Analysis of Chlamydia Trachomatis Oculotropic and Genitotropic Strains. Infect Immun 2005, 73, 6407–6418, doi:10.1128/IAI.73.10.6407-6418.2005.

51. Andersson, P.; Harris, S.R.; Smith, H.M.B.S.; Hadfield, J.; O’Neill, C.; Cutcliffe, L.T.; Douglas, F.P.; Asche, L.V.; Mathews, J.D.; Hutton, S.I.;, et al. Chlamydia Trachomatis from Australian Aboriginal People with Trachoma Are Polyphyletic Composed of Multiple Distinctive Lineages. Nat Commun 2016, 7, 10688, doi:10.1038/ncomms10688.

52. Seth-Smith, H.M.B.; Harris, S.R.; Persson, K.; Marsh, P.; Barron, A.; Bignell, A.; Bjartling, C.; Clark, L.; Cutcliffe, L.T.; Lambden, P.R.;, et al. Co-Evolution of Genomes and Plasmids within Chlamydia Trachomatis and the Emergence in Sweden of a New Variant Strain. BMC Genomics 2009, 10, 239, doi:10.1186/1471-2164-10-239.

53. Jones, C.A.; Hadfield, J.; Thomson, N.R.; Cleary, D.W.; Marsh, P.; Clarke, I.N.; O’Neill, C.E. The Nature and Extent of Plasmid Variation in Chlamydia Trachomatis. Microorganisms 2020, 8, 373, doi:10.3390/microorganisms8030373.

54. Fisher, D.J.; Fernández, R.E.; Maurelli, A.T. Chlamydia Trachomatis Transports NAD via the Npt1 ATP/ADP Translocase. J Bacteriol 2013, 195, 3381–3386, doi:10.1128/JB.00433-13.

55. Rucks, E.A. Type III Secretion in Chlamydia. Microbiol Mol Biol Rev 2023, 87, e0003423, doi:10.1128/mmbr.00034-23.

56. Tanaka, K.J.; Song, S.; Mason, K.; Pinkett, H.W. Selective Substrate Uptake: The Role of ATP-Binding Cassette (ABC) Importers in Pathogenesis. Biochim Biophys Acta Biomembr 2018, 1860, 868–877, doi:10.1016/j.bbamem.2017.08.011.

57. Kari, L.; Whitmire, W.M.; Carlson, J.H.; Crane, D.D.; Reveneau, N.; Nelson, D.E.; Mabey, D.C.; Bailey, R.L.; Holland, M.J.; McClarty, G.;, et al. Pathogenic Diversity among Chlamydia Trachomatis Ocular Strains in Nonhuman Primates Is Affected by Subtle Genomic Variations. J Infect Dis 2008, 197, 449–456, doi:10.1086/525285.

58. Braun, C.; Hegemann, J.H.; Mölleken, K. Insights Into a Chlamydia Pneumoniae-Specific Gene Cluster of Membrane Binding Proteins. Front Cell Infect Microbiol 2020, 10, 565808, doi:10.3389/fcimb.2020.565808.

59. Weber, M.M.; Bauler, L.D.; Lam, J.; Hackstadt, T. Expression and Localization of Predicted Inclusion Membrane Proteins in Chlamydia Trachomatis. Infect Immun 2015, 83, 4710–4718, doi:10.1128/IAI.01075-15.

60. Weber, M.M.; Noriea, N.F.; Bauler, L.D.; Lam, J.L.; Sager, J.; Wesolowski, J.; Paumet, F.; Hackstadt, T. A Functional Core of IncA Is Required for Chlamydia Trachomatis Inclusion Fusion. J Bacteriol 2016, 198, 1347–1355, doi:10.1128/JB.00933-15.

61. Thomson, N.R.; Holden, M.T.G.; Carder, C.; Lennard, N.; Lockey, S.J.; Marsh, P.; Skipp, P.; O’Connor, C.D.; Goodhead, I.; Norbertzcak, H.;, et al. Chlamydia Trachomatis: Genome Sequence Analysis of Lymphogranuloma Venereum Isolates. Genome Res 2008, 18, 161–171, doi:10.1101/gr.7020108.

62. Saini, A.; Mapolelo, D.T.; Chahal, H.K.; Johnson, M.K.; Outten, F.W. SufD and SufC ATPase Activity Are Required for Iron Acquisition during in Vivo Fe-S Cluster Formation on SufB. Biochemistry 2010, 49, 9402–9412, doi:10.1021/bi1011546.

63. Hamaoui, D.; Cossé, M.M.; Mohan, J.; Lystad, A.H.; Wollert, T.; Subtil, A. The Chlamydia Effector CT622/TaiP Targets a Nonautophagy Related Function of ATG16L1. Proc Natl Acad Sci U S A 2020, 117, 26784–26794, doi:10.1073/pnas.2005389117.

64. Auer, D.; Hügelschäffer, S.D.; Fischer, A.B.; Rudel, T. The Chlamydial Deubiquitinase Cdu1 Supports Recruitment of Golgi Vesicles to the Inclusion. Cell Microbiol 2020, 22, e13136, doi:10.1111/cmi.13136.

65. Hausman, J.M.; Kenny, S.; Iyer, S.; Babar, A.; Qiu, J.; Fu, J.; Luo, Z.-Q.; Das, C. The Two Deubiquitinating Enzymes from Chlamydia Trachomatis Have Distinct Ubiquitin Recognition Properties. Biochemistry 2020, 59, 1604–1617, doi:10.1021/acs.biochem.9b01107.

66. Huang, Y.; Wurihan, W.; Lu, B.; Zou, Y.; Wang, Y.; Weldon, K.; Fondell, J.D.; Lai, Z.; Wu, X.; Fan, H. Robust Heat Shock Response in Chlamydia Lacking a Typical Heat Shock Sigma Factor. Front Microbiol 2021, 12, 812448, doi:10.3389/fmicb.2021.812448.

67. Benamri, I.; Azzouzi, M.; Moussa, A.; Radouani, F. An in Silico Analysis of RpoB Mutations to Affect Chlamydia Trachomatis Sensitivity to Rifamycin. J Genet Eng Biotechnol 2022, 20, 146, doi:10.1186/s43141-022-00428-y.

68. Gorbalenya, A.E.; Koonin, E. V; Donchenko, A.P.; Blinov, V.M. Two Related Superfamilies of Putative Helicases Involved in Replication, Recombination, Repair and Expression of DNA and RNA Genomes. Nucleic Acids Res 1989, 17, 4713–4730, doi:10.1093/nar/17.12.4713.

69. Soules, K.R.; LaBrie, S.D.; May, B.H.; Hefty, P.S. Sigma 54-Regulated Transcription Is Associated with Membrane Reorganization and Type III Secretion Effectors during Conversion to Infectious Forms of Chlamydia Trachomatis. mBio 2020, 11, doi:10.1128/mBio.01725-20.

70. Ilic, S.; Cohen, S.; Singh, M.; Tam, B.; Dayan, A.; Akabayov, B. DnaG Primase-A Target for the Development of Novel Antibacterial Agents. Antibiotics (Basel) 2018, 7, doi:10.3390/antibiotics7030072.

71. Wang, L.; Hou, Y.; Yuan, H.; Chen, H. The Role of Tryptophan in Chlamydia Trachomatis Persistence. Front Cell Infect Microbiol 2022, 12, 931653, doi:10.3389/fcimb.2022.931653.

72. Wylie, J.L.; Berry, J.D.; McClarty, G. Chlamydia Trachomatis CTP Synthetase: Molecular Characterization and Developmental Regulation of Expression. Mol Microbiol 1996, 22, 631–642, doi:10.1046/j.1365-2958.1996.d01-1717.x.

73. Lorca, G.L.; Barabote, R.D.; Zlotopolski, V.; Tran, C.; Winnen, B.; Hvorup, R.N.; Stonestrom, A.J.; Nguyen, E.; Huang, L.-W.; Kim, D.S.;, et al. Transport Capabilities of Eleven Gram-Positive Bacteria: Comparative Genomic Analyses. Biochim Biophys Acta 2007, 1768, 1342–1366, doi:10.1016/j.bbamem.2007.02.007.

74. Griffin, J.; Roshick, C.; Iliffe-Lee, E.; McClarty, G. Catalytic Mechanism of Chlamydia Trachomatis Flavin-Dependent Thymidylate Synthase. J Biol Chem 2005, 280, 5456– 5467, doi:10.1074/jbc.M412415200.

75. Kari, L.; Southern, T.R.; Downey, C.J.; Watkins, H.S.; Randall, L.B.; Taylor, L.D.; Sturdevant, G.L.; Whitmire, W.M.; Caldwell, H.D. Chlamydia Trachomatis Polymorphic Membrane Protein D Is a Virulence Factor Involved in Early Host-Cell Interactions. Infect Immun 2014, 82, 2756–2762, doi:10.1128/IAI.01686-14.

76. Yang, M.; Rajeeve, K.; Rudel, T.; Dandekar, T. Comprehensive Flux Modeling of Chlamydia Trachomatis Proteome and QRT-PCR Data Indicate Biphasic Metabolic Differences Between Elementary Bodies and Reticulate Bodies During Infection. Front Microbiol 2019, 10, 2350, doi:10.3389/fmicb.2019.02350.

77. Binet, R.; Fernandez, R.E.; Fisher, D.J.; Maurelli, A.T. Identification and Characterization of the Chlamydia Trachomatis L2 S-Adenosylmethionine Transporter. mBio 2011, 2, e00051–11, doi:10.1128/mBio.00051-11.

78. Stephens, R.S.; Kalman, S.; Lammel, C.; Fan, J.; Marathe, R.; Aravind, L.; Mitchell, W.; Olinger, L.; Tatusov, R.L.; Zhao, Q.;, et al. Genome Sequence of an Obligate Intracellular Pathogen of Humans: Chlamydia Trachomatis. Science 1998, 282, 754– 759, doi:10.1126/science.282.5389.754.

79. Taylor, L.D.; Nelson, D.E.; Dorward, D.W.; Whitmire, W.M.; Caldwell, H.D. Biological Characterization of Chlamydia Trachomatis Plasticity Zone MACPF Domain Family Protein CT153. Infect Immun 2010, 78, 2691–2699, doi:10.1128/IAI.01455-09.

80. Belland, R.J.; Zhong, G.; Crane, D.D.; Hogan, D.; Sturdevant, D.; Sharma, J.; Beatty, W.L.; Caldwell, H.D. Genomic Transcriptional Profiling of the Developmental Cycle of Chlamydia Trachomatis. Proc Natl Acad Sci U S A 2003, 100, 8478–8483, doi:10.1073/pnas.1331135100.

81. Macleod, C.K.; Butcher, R.; Mudaliar, U.; Natutusau, K.; Pavluck, A.L.; Willis, R.; Alexander, N.; Mabey, D.C.; Cikamatana, L.; Kama, M.;, et al. Low Prevalence of Ocular Chlamydia Trachomatis Infection and Active Trachoma in the Western Division of Fiji. PLoS Negl Trop Dis 2016, 10, e0004798, doi:10.1371/journal.pntd.0004798.

82. Harris, S.R.; Clarke, I.N.; Seth-Smith, H.M.B.; Solomon, A.W.; Cutcliffe, L.T.; Marsh, P.; Skilton, R.J.; Holland, M.J.; Mabey, D.; Peeling, R.W.;, et al. Whole Genome Analysis of Diverse Chlamydia Trachomatis Strains Identifies Phylogenetic Relationships Masked by Current Clinical Typing. Nat Genet 2012, 44, 413–S1, doi:10.1038/ng.2214.

83. Jeffrey, B.M.; Suchland, R.J.; Quinn, K.L.; Davidson, J.R.; Stamm, W.E.; Rockey, D.D. Genome Sequencing of Recent Clinical Chlamydia Trachomatis Strains Identifies Loci Associated with Tissue Tropism and Regions of Apparent Recombination. Infect Immun 2010, 78, 2544–2553, doi:10.1128/IAI.01324-09.

84. Versteeg, B.; Bruisten, S.M.; Pannekoek, Y.; Jolley, K.A.; Maiden, M.C.J.; van der Ende, A.; Harrison, O.B. Genomic Analyses of the Chlamydia Trachomatis Core Genome Show an Association between Chromosomal Genome, Plasmid Type and Disease. BMC Genomics 2018, 19, 130, doi:10.1186/s12864-018-4522-3.

85. Seth-Smith, H.M.B.; Bénard, A.; Bruisten, S.M.; Versteeg, B.; Herrmann, B.; Kok, J.; Carter, I.; Peuchant, O.; Bébéar, C.; Lewis, D.A.;, et al. Ongoing Evolution of Chlamydia Trachomatis Lymphogranuloma Venereum: Exploring the Genomic Diversity of Circulating Strains. Microb Genom 2021, 7, doi:10.1099/mgen.0.000599.

86. Read, T.D.; Joseph, S.J.; Didelot, X.; Liang, B.; Patel, L.; Dean, D. Comparative Analysis of Chlamydia Psittaci Genomes Reveals the Recent Emergence of a Pathogenic Lineage with a Broad Host Range. mBio 2013, 4, doi:10.1128/mBio.00604-12.

87. Williams, M.J.; Zapata, L.; Werner, B.; Barnes, C.P.; Sottoriva, A.; Graham, T.A. Measuring the Distribution of Fitness Effects in Somatic Evolution by Combining Clonal Dynamics with DN/DS Ratios. Elife 2020, 9, doi:10.7554/eLife.48714.

88. Wolf, J.B.W.; Künstner, A.; Nam, K.; Jakobsson, M.; Ellegren, H. Nonlinear Dynamics of Nonsynonymous (DN) and Synonymous (DS) Substitution Rates Affects Inference of Selection. Genome Biol Evol 2009, 1, 308–319, doi:10.1093/gbe/evp030.

89. Kryazhimskiy, S.; Plotkin, J.B. The Population Genetics of DN/DS. PLoS Genet 2008, 4, e1000304, doi:10.1371/journal.pgen.1000304.

90. Borges, V.; Gomes, J.P. Deep Comparative Genomics among Chlamydia Trachomatis Lymphogranuloma Venereum Isolates Highlights Genes Potentially Involved in Pathoadaptation. Infect Genet Evol 2015, 32, 74–88, doi:10.1016/j.meegid.2015.02.026.

91. Kari, L.; Whitmire, W.M.; Carlson, J.H.; Crane, D.D.; Reveneau, N.; Nelson, D.E.; Mabey, D.C.W.; Bailey, R.L.; Holland, M.J.; McClarty, G.;, et al. Pathogenic Diversity among Chlamydia Trachomatis Ocular Strains in Nonhuman Primates Is Affected by Subtle Genomic Variations. J Infect Dis 2008, 197, 449–456, doi:10.1086/525285.

92. Wang, L.; Hou, Y.; Yuan, H.; Chen, H. The Role of Tryptophan in Chlamydia Trachomatis Persistence. Front Cell Infect Microbiol 2022, 12, 931653, doi:10.3389/fcimb.2022.931653.

93. Sigalova, O.M.; Chaplin, A. V; Bochkareva, O.O.; Shelyakin, P. V; Filaretov, V.A.; Akkuratov, E.E.; Burskaia, V.; Gelfand, M.S. Chlamydia Pan-Genomic Analysis Reveals Balance between Host Adaptation and Selective Pressure to Genome Reduction. BMC Genomics 2019, 20, 710, doi:10.1186/s12864-019-6059-5.

94. Caldwell, H.D.; Wood, H.; Crane, D.; Bailey, R.; Jones, R.B.; Mabey, D.; Maclean, I.; Mohammed, Z.; Peeling, R.; Roshick, C.;, et al. Polymorphisms in Chlamydia Trachomatis Tryptophan Synthase Genes Differentiate between Genital and Ocular Isolates. Journal of Clinical Investigation 2003, 111, 1757–1769, doi:10.1172/JCI17993.

95. Smelov, V.; Vrbanac, A.; van Ess, E.F.; Noz, M.P.; Wan, R.; Eklund, C.; Morgan, T.; Shrier, L.A.; Sanders, B.; Dillner, J.;, et al. Chlamydia Trachomatis Strain Types Have Diversified Regionally and Globally with Evidence for Recombination across Geographic Divides. Front Microbiol 2017, 8, doi:10.3389/fmicb.2017.02195.

96. Joseph, S.J.; Bommana, S.; Ziklo, N.; Kama, M.; Dean, D.; Read, T.D. Patterns of Within-Host Spread of Chlamydia Trachomatis between Vagina, Endocervix and Rectum Revealed by Comparative Genomic Analysis. Front Microbiol 2023, 14, 1154664, doi:10.3389/fmicb.2023.1154664.

97. Bom, R.J.M.; Christerson, L.; Schim van der Loeff, M.F.; Coutinho, R.A.; Herrmann, B.; Bruisten, S.M. Evaluation of High-Resolution Typing Methods for Chlamydia Trachomatis in Samples from Heterosexual Couples. J Clin Microbiol 2011, 49, 2844– 2853, doi:10.1128/JCM.00128-11.

98. Chidambaram, J.D.; Lee, D.C.; Porco, T.C.; Lietman, T.M. Mass Antibiotics for Trachoma and the Allee Effect. Lancet Infect Dis 2005, 5, 194–196, doi:10.1016/S1473-3099(05)70032-3.

