## Supplementary material for "Genomic insights into local scale evolution of ocular *Chlamydia trachomatis* strains within and between individuals in Gambian trachoma-endemic villages": Data S1

Data S1: Supporting Information for

### Genomic insights into local scale evolution of ocular *Chlamydia trachomatis* strains within and between individuals in Gambian trachoma-endemic villages

Ehsan Ghasemian[1*], Nkoyo Faal[2], Harry Pickering[1], Ansumana Sillah[3], Judith Breuer[4], Robin L. Bailey[1], David Mabey[1], Martin J. Holland[1]

[1] Department of Clinical Research, London School of Hygiene & Tropical Medicine, London, United Kingdom

[2] Medical Research Council Unit The Gambia at London School of Hygiene and Tropical Medicine, Banjul, The Gambia

[3] National Eye Health Programme, Ministry of Health, Kanifing, The Gambia

[4] Division of Infection and Immunity, University College London, London, United Kingdom

#### *Chlamydia trachomatis* strains B/HAR36 and B/Tunis864

The London School of Hygiene & Tropical Medicine (LSHTM) trachoma group obtained live stocks and genomic DNA from *Chlamydia trachomatis* (Ct) strains A/2497, A/2497P-, B/HAR36 and C/TW3 via Harlan Caldwell at the Laboratory of Intracellular Parasites, Rocky Mountain Laboratories, National Institute of Allergy and Infectious Diseases, National Institutes of Health, Hamilton, Montana. Comparative genomic investigations on these strains were published by Kari *et al*. [1]. LSHTM provided the Wellcome Trust Sanger Institute (WTSI) genomic DNA from A/2497, B/HAR36 and C/TW3, which underwent full genome sequencing by Next Generation Sequencing and the sequences deposited in the European Nucleotide Archive (ENA) under the following accession numbers (CP002401, ERR189736, and CP006945, respectively). These sequences were included in analyses by Andersson *et al*. [2] and Hadfield *et al*. [3]. At the start of the EU Horizon-2020 funded TracVac project (2017 – 2022, Grant agreement code: 733373) B/HAR36 was selected for large scale culture to represent serovar B genotypes as part of *in vitro* and *in vivo* vaccine studies. Subsequently DNA extracted from these B/HAR36 cultures were subjected to *omp*A chain-termination sequencing (Sanger sequencing). B/HAR36 labelled *omp*A sequence was found to be a 100% match for NCBI accession DQ064280 *omp*A from B/Tunis864 with a lower identity (99%) to NCBI accession DQ064297.1 *omp*A sequence from B/HAR36. *omp*A sequence from whole genome sequence ERR189736 was extracted and was a match for B/Tunis864 (DQ064280). We therefore, obtained original stocks of B/HAR36 and B/Tunis864 from Julius Schachter and Jeanne Moncada at the World Health Organization Collaborating Centre for Reference and Research on Trachoma and Other Chlamydial Infections, Francis I Proctor Foundation for research in Ophthalmology, University of California, San Francisco (UCSF) and subjected these to Whole-Genome Sequencing along with ERR189736 DNA originally supplied to WTSI. We found that the whole genome sequence initially deposited to the ENA as strain

B/HAR36 (ERR189736) was a closer match with the new genome sequence of strain B/Tunis864 with three variations between the sequences. The new B/Tunis864 genome sequence, is now deposited under the ENA accession number ERR12253485. B/HAR36 original stock obtained from UCSF was distinct from B/Tunis864 at the whole genome level with 1,407 variations and the extracted *omp*A sequence was a match (100%) for NCBI DQ064297.1 *omp*A (B/HAR36 *omp*A); the new genome sequence of strain B/HAR36 is now deposited as ERR12253486. Since the full genome sequence ERR189736 initially labelled as B/HAR36 and B/Tunis864 full genome sequence (ERR12253485) represent the same strain genome, the observation that ERR189736 carried B/Tunis864 *omp*A is not explained by *omp*A recombination during laboratory culture as at the time of culture B/HAR36 was not available in the LSHTM laboratories or at the Statens serum Institute, Copenhagen, Denmark where bulk culture was performed as part of TracVac. We therefore infer that there was a labelling mistake on stock that was sequenced to generate ERR189736; correction of metadata for this record is underway.

**References**


1. Kari, L.; Whitmire, W.M.; Crane, D.D.; Reveneau, N.; Carlson, J.H.; Goheen, M.M.; Peterson, E.M.; Pal, S.; de la Maza, L.M.; Caldwell, H.D. Chlamydia Trachomatis Native Major Outer Membrane Protein Induces Partial Protection in Nonhuman Primates: Implication for a Trachoma Transmission-Blocking Vaccine. *The Journal of Immunology* **2009**, *182*, 8063, doi:10.4049/jimmunol.0804375.
2. Andersson, P.; Harris, S.R.; Smith, H.M.B.S.; Hadfield, J.; O'Neill, C.; Cutcliffe, L.T.; Douglas, F.P.; Asche, L.V.; Mathews, J.D.; Hutton, S.I.; et al. Chlamydia Trachomatis from Australian Aboriginal People with Trachoma Are Polyphyletic Composed of

Multiple Distinctive Lineages. *Nat Commun* **2016**, *7*, 10688, doi:10.1038/ncomms10688.

3. Hadfield, J.; Harris, S.R.; Seth-Smith, H.M.B.; Parmar, S.; Andersson, P.; Giffard, P.M.; Schachter, J.; Moncada, J.; Ellison, L.; Vaulet, M.L.G.; et al. Comprehensive Global Genome Dynamics of Chlamydia Trachomatis Show Ancient Diversification Followed by Contemporary Mixing and Recent Lineage Expansion. *Genome Res* **2017**, *27*, 1220–1229, doi:10.1101/gr.212647.116.
