## Supplementary material for "Genomic insights into local scale evolution of ocular *Chlamydia trachomatis* strains within and between individuals in Gambian trachoma-endemic villages": Data S2

Data S2: Supporting Information for

### Genomic insights into local scale evolution of ocular *Chlamydia trachomatis* strains within and between individuals in Gambian trachoma-endemic villages

Ehsan Ghasemian[1*], Nkoyo Faal[2], Harry Pickering[1], Ansumana Sillah[3], Judith Breuer[4], Robin L. Bailey[1], David Mabey[1], Martin J. Holland[1]

[1] Department of Clinical Research, London School of Hygiene & Tropical Medicine, London, United Kingdom

[2] Medical Research Council Unit The Gambia at London School of Hygiene and Tropical Medicine, Banjul, The Gambia

[3] National Eye Health Programme, Ministry of Health, Kanifing, The Gambia

[4] Division of Infection and Immunity, University College London, London, United Kingdom

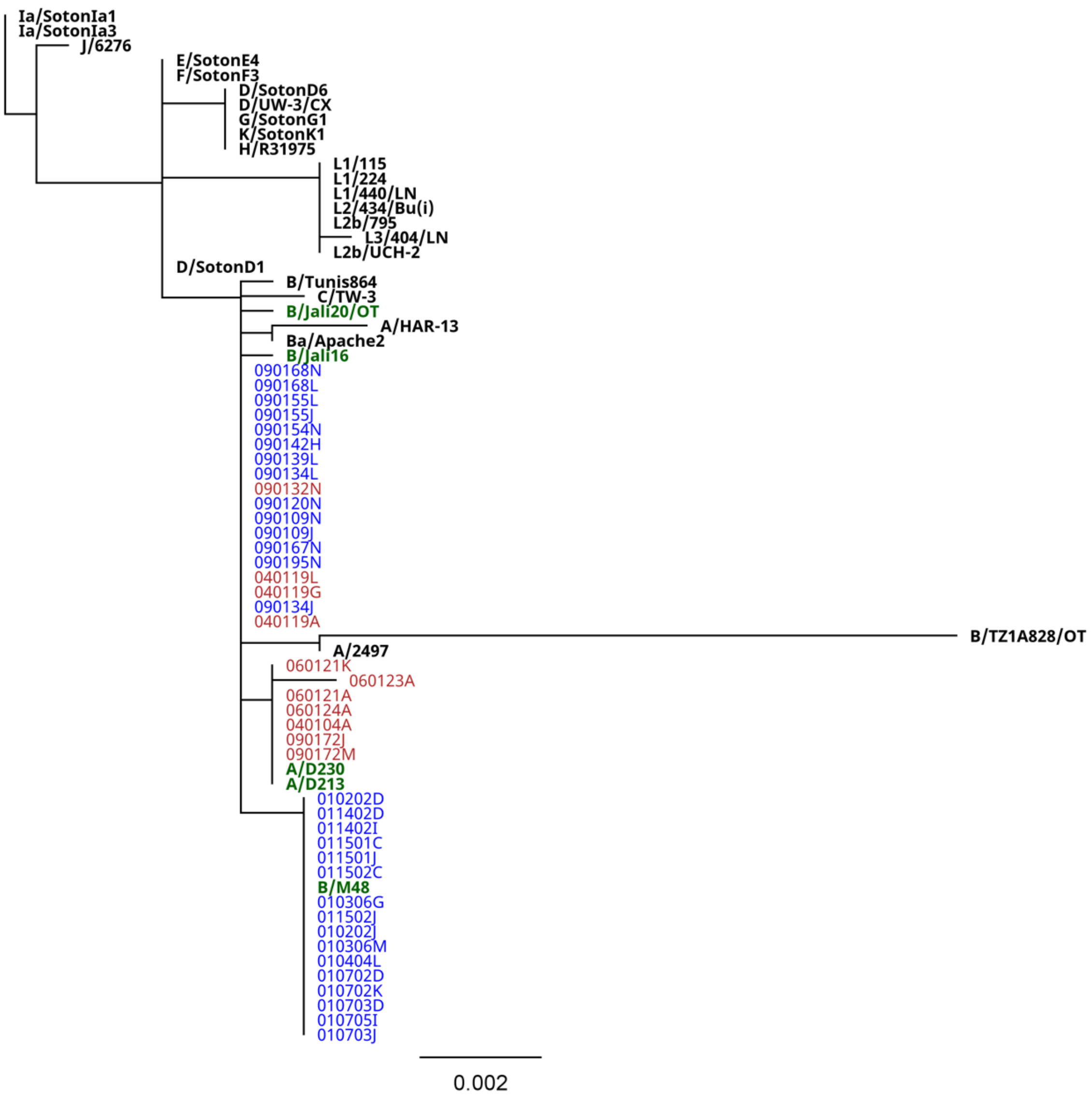

**Fig. S1.** Phylogenetic analysis of *Chlamydia trachomatis trp*AB genes. The phylogenetic tree encompasses 41 *C. trachomatis* (Ct) positive samples obtained from The Gambia, alongside 29 Ct reference strains. Sequence alignments were generated using MAFFT, and maximum likelihood phylogenies of these aligned sequences were estimated employing the General Time Reversible (GTR) model of evolution with 1000 bootstrap replicates, facilitated by PhyML.

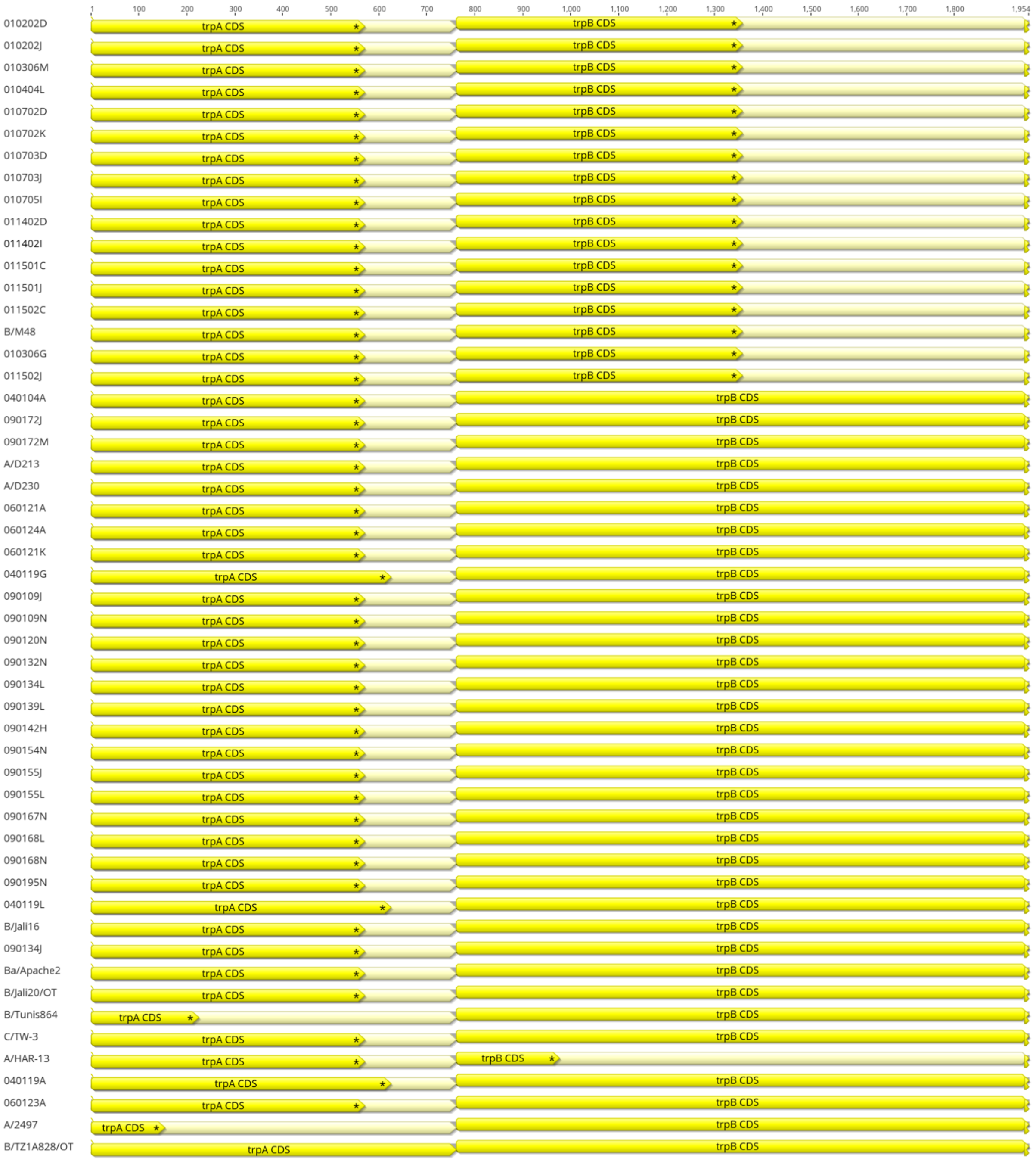

**Fig. S2.** Comparative alignment of *Chlamydia trachomatis trp*AB Genes. Alignment of *trp*AB genes from *C. trachomatis* (Ct) sequences sourced from the Gambia and Ct ocular reference strains was conducted using MAFFT (version v7.490). The asterisk in the annotation indicates the termination of the coding sequence.

**Table S1.** Baseline demographics, trachoma grades, CT * omp*A type, and *omc*B load of study participants and sequences quality information.

| | Sample ID | Sex | Age | Trachoma grade | pORF2 load | *omc*B load | *omp*A genotype | Plasmid genotype | Number of raw read pairs | Number of read pairs after adaptor trimming, qualiy score filtering (>=20), and merging | Per base sequence quality | Average quality per read | % GC | % per base N content | Mean length | Number of read paired classified as *Chlamydia trachomatis* | Mean coverag of *Chlamydia trachomatis* reference genome | Std Dev (mean coverag of *Chlamydia trachomatis* reference genome) | Ref-seq (%) | Confidence mean |
|---|---|---|---|---|---|---|---|---|---|---|---|---|---|---|---|---|---|---|---|---|
| **Village 1** | | | | | | | | | | | | | | | | | | | | |
| | 010202_D | F | 5 | TF | 79.4 | 12.5 | B | B | 3,607,364 | 1,413,099 | >28 | 38 | 40 | 0 | 153 | 122,173 | 21.3 | 14 | 99.4 | 37.5 |
| | 010202_J | F | 5 | TF | 121.5 | 17.1 | B | B | 3,608,774 | 1,191,509 | >28 | 38 | 43 | 0 | 151 | 356,920 | 53.6 | 28 | 99.6 | 37.6 |
| | 010306_G | F | 5 | TF | 22992.1 | 3071.4 | B | B | 4,041,512 | 1,374,631 | >28 | 38 | 41 | 0 | 170 | 914,801 | 142 | 54.5 | 99.6 | 37.5 |
| | 010306_M | F | 5 | TF | 1107.6 | 184.6 | B | B | 3,859,352 | 1,480,843 | >28 | 38 | 41 | 0 | 162 | 238,390 | 41.7 | 25.5 | 99.6 | 37.6 |
| | 010404_L | F | 10 | TF | 679 | 37.3 | B | B | 3,495,140 | 1,114,702 | >28 | 37 | 41 | 0 | 163 | 545,362 | 85.7 | 35.3 | 99.6 | 37.4 |
| | 010702_D | F | 7 | TF | 318.6 | 54.4 | B | B | 2,476,052 | 677,749 | >28 | 38 | 41 | 0 | 168 | 396,714 | 65.8 | 24.8 | 99.6 | 37 |
| | 010702_K | F | 7 | TF | 312.4 | 58.5 | B | B | 3,622,540 | 1,328,892 | >28 | 37 | 43 | 0 | 170 | 452,429 | 80.9 | 48.3 | 99.6 | 37.5 |
| | 010703_D | F | 4 | TF | 9736.2 | 1388.1 | B | B | 3,304,646 | 937,981 | >28 | 37 | 41 | 0 | 172 | 636,957 | 105.2 | 40.6 | 99.6 | 37 |
| | 010703_J | F | 4 | TF | 4619.7 | 837.1 | B | B | 2,511,736 | 724,754 | >28 | 37 | 41 | 0 | 163 | 483,233 | 80.7 | 31.1 | 99.6 | 37 |
| | 010705_I | M | 10 | Normal | 525.8 | 68.8 | B | B | 3,151,364 | 1,022,956 | >28 | 37 | 41 | 0 | 172 | 591,994 | 94.2 | 36.1 | 99.6 | 37.3 |
| | 011402_D | F | 10 | TF | 17460 | 3063.6 | B | B | 4,590,522 | 1,281,634 | >28 | 37 | 41 | 0 | 172 | 876,389 | 146.5 | 54.3 | 99.6 | 36.9 |
| | 011402_I | F | 10 | TF | 5074.9 | 857.8 | B | B | 4,529,892 | 1,279,478 | >28 | 37 | 41 | 0 | 172 | 887,090 | 148.7 | 54.1 | 99.6 | 37 |
| | 011501_C | F | 5 | TF | 2503.1 | 303.5 | B | B | 3,573,360 | 1,156,467 | >28 | 37 | 41 | 0 | 163 | 743,013 | 117.1 | 45.8 | 99.6 | 37.4 |
| | 011501_J | F | 5 | TF | 6200.4 | 2798.9 | B | B | 6,688,138 | 1,913,661 | >28 | 37 | 41 | 0 | 172 | 1,334,258 | 222.4 | 80.2 | 99.6 | 37 |
| | 011502_C | F | 4 | TF | 1214.3 | 279.8 | B | B | 3,913,356 | 1,463,596 | >28 | 38 | 42 | 0 | 179 | 1,196,075 | 210.1 | 115.6 | 99.6 | 37.6 |
| | 011502_J | F | 4 | TF | 767.8 | 106.4 | B | B | 3,076,780 | 874,013 | >28 | 37 | 42 | 0 | 169 | 432,783 | 71.7 | 29.3 | 99.6 | 37.1 |
| **Village 4** | | | | | | | | | | | | | | | | | | | | |
| | 040104_A | M | 6 | TF | 1646 | 249.3 | A | A | 2,337,432 | 611,793 | >28 | 37 | 42 | 0 | 173 | 347,444 | 58.6 | 24.1 | 99.6 | 36.9 |
| | 040119_A | M | 13 | TF | 137.8 | 27 | A | A | 2,457,774 | 637,256 | >28 | 37 | 43 | 0 | 169 | 246,632 | 41.4 | 18.2 | 99.5 | 36.9 |
| | 040119_G | M | 13 | TF | 250 | 31.8 | A | A | 3,684,418 | 1,313,457 | >28 | 38 | 44 | 0 | 166 | 406,104 | 68 | 32.3 | 99.5 | 37.8 |
| | 040119_L | M | 13 | TF | 157.1 | 22.6 | A | A | 3,259,084 | 1,031,894 | >28 | 38 | 43 | 0 | 156 | 327,882 | 51.6 | 22.8 | 99.5 | 37.4 |
| **Village 6** | | | | | | | | | | | | | | | | | | | | |
| | 060121_A | M | 8 | TF | 7516.9 | 653.7 | A | A | 3,431,668 | 1,149,974 | >28 | 38 | 41 | 0 | 161 | 723,676 | 113.9 | 47.2 | 98.1 | 37.4 |
| | 060121_K | M | 8 | Normal | 761 | 63.6 | A | A | 2,517,680 | 668,524 | >28 | 37 | 42 | 0 | 166 | 273,864 | 44.4 | 21.4 | 98.1 | 37.2 |
| | 060123_A | M | 7 | TF | 1830.2 | 216 | A | A | 4,191,512 | 1,571,119 | >28 | 38 | 41 | 0 | 149 | 908,894 | 132.6 | 64 | 98 | 37.9 |
| | 060124_A | F | 8 | TF | 287.2 | 24.7 | A | A | 3,342,888 | 904,124 | >28 | 37 | 42 | 0 | 167 | 327,738 | 54.3 | 24 | 98.1 | 37.1 |
| **Village 9** | | | | | | | | | | | | | | | | | | | | |
| | 090109_J | M | 15 | TF | 2556.8 | 639.2 | B | B | 4,799,546 | 2,171,961 | >28 | 40 | 42 | 0 | 199 | 1,332,150 | 270.9 | 154.2 | 99.6 | 40 |
| | 090109_N | M | 15 | Normal | 997.5 | 176.8 | B | B | 3,287,146 | 1,558,956 | >28 | 40 | 43 | 0 | 203 | 901,921 | 178 | 67 | 99.6 | 40.3 |
| | 090120_N | M | 9 | Normal | 342.5 | 69.3 | B | B | 3,638,672 | 1,748,766 | >28 | 41 | 42 | 0 | 170 | 951,485 | 165.4 | 64.7 | 99.6 | 40.4 |
| | 090132_N | M | 13 | TF | 147.5 | 29.5 | A | A | 3,881,366 | 1,804,625 | >28 | 40 | 44 | 0 | 205 | 708,070 | 144.3 | 56.9 | 99.6 | 40.2 |
| | 090134_J | M | 9 | TF | 235 | 44.9 | B | B | 3,785,094 | 1,446,284 | >28 | 38 | 43 | 0 | 157 | 108,097 | 19.1 | 17.1 | 96.2 | 37.5 |
| | 090134_L | M | 9 | TF | 802.1 | 183.8 | B | B | 2,389,658 | 647,184 | >28 | 37 | 42 | 0 | 169 | 325,673 | 54.2 | 22 | 99.6 | 37 |
| | 090139_L | M | 9 | TF | 182.5 | 34.5 | B | B | 3,614,836 | 1,695,248 | >28 | 40 | 43 | 0 | 204 | 837,136 | 164.1 | 63.2 | 99.6 | 40.2 |
| | 090142_H | M | 13 | TF | 116.4 | 27 | B | A | 3,377,450 | 1,158,843 | >28 | 38 | 43 | 0 | 174 | 406,393 | 70.5 | 30 | 99.6 | 37.6 |
| | 090154_N | M | 11 | Normal | 307.5 | 69 | B | B | 3,510,798 | 1,642,701 | >28 | 40 | 43 | 0 | 214 | 891,817 | 187.1 | 69.4 | 99.6 | 40.1 |
| | 090155_J | M | 9 | TF | 192.1 | 41.6 | B | B | 3,359,078 | 1,289,653 | >28 | 38 | 41 | 0 | 164 | 400,653 | 70.3 | 43.2 | 99.6 | 37.6 |
| | 090155_L | M | 9 | TF | 2424.5 | 553.1 | B | B | 2,692,100 | 765,780 | >28 | 37 | 42 | 0 | 171 | 507,780 | 84.7 | 33 | 99.6 | 37 |
| | 090167_N | M | 9 | Normal | 153.7 | 28.8 | B | B | 3,276,578 | 1,519,348 | >28 | 40 | 44 | 0 | 205 | 495,844 | 102.1 | 44.4 | 99.6 | 40.1 |
| | 090168_L | M | 10 | TF | 1780 | 500.8 | B | B | 3,649,316 | 1,225,599 | >28 | 38 | 42 | 0 | 181 | 683,884 | 121.7 | 43.9 | 99.6 | 37.4 |
| | 090168_N | M | 10 | Normal | 297.9 | 55.6 | B | B | 3,598,198 | 1,717,357 | >28 | 40 | 42 | 0 | 195 | 1,163,730 | 227.5 | 81.3 | 99.6 | 40.2 |
| | 090172_J | M | 11 | TF | 5449.3 | 1052 | A | A | 3,888,716 | 1,324,270 | >28 | 38 | 41 | 0 | 180 | 763,165 | 134.6 | 49 | 99.6 | 37.4 |
| | 090172_M | M | 11 | TF | 4311.9 | 815.2 | A | A | 3,442,574 | 1,051,627 | >28 | 37 | 41 | 0 | 168 | 697,247 | 113.9 | 43.2 | 99.6 | 37.2 |
| | 090195_N | M | 8 | TF | 68.5 | 37.5 | B | B | 3,698,022 | 1,765,573 | >28 | 40 | 42 | 0 | 207 | 1,260,658 | 255.5 | 94.6 | 99.6 | 40.2 |

**Table S2.** List of *Chlamydia trachomatis* reference strains used in this study and metadata for the sequences.

| Isolate name | Lineage | Genotype | Country | Year | Source | Genome-ERR | Genome-ERS | Genome-accession number | Genome-reference | Plasmid-ERR | Plasmid-ERS | Plasmid-accession number | Plasmid- reference |
|---|---|---|---|---|---|---|---|---|---|---|---|---|---|
| A/2497 | ocular | A | Tanzania | 2000 | ocular | - | - | FM872306 | Harris et al. 2012 | - | - | NC_020550 | Clarke and Seth-Smith. 2008 |
| A/D213 | ocular | A | Gambia | 2001 | ocular | ERR175652 | ERS177838 | - | Andersson et al. 2016 | ERR175652 | ERS177838 | - | - |
| A/D230 | ocular | A | Gambia | 2001 | ocular | ERR111554 | ERS075177 | - | Hadfield et al. 2017 | ERR111554 | ERS075177 | - | - |
| A/HAR13 | ocular | A | Egypt | 1958 | conjunctiva | | - | CP000051 | Carlson 2005 | - | - | NC_007430 | Carlson et al., 2005 |
| B/Jali16 | ocular | B | Gambia | 1985 | ocular | ERR189738 | ERS153015 | - | Hadfield et al. 2017 | ERR189738 | ERS153015 | - | - |
| B/Jali20 | ocular | B | Gambia | 1985 | ocular | - | - | FM872308 | Seth-Smith et al 2009 | - | - | NC_012629 | Seth-Smith et al. 2009 |
| B/TZ1A828 | ocular | B | Tanzania | 1998 | ocular | - | - | FM872307 | Seth-Smith et al 2009 | - | - | FM865437 | Seth-Smith et al. 2009 |
| B/M48 | ocular | B | Gambia | 2007 | ocular | ERR175631 | ERS177817 | - | Hadfield et al. 2017 | ERR175631 | ERS177817 | - | - |
| B/Tunis864 | ocular | B | Tunisia | 1976 | ocular | ERR12253485 | ERS16770978 | - | This study | ERR12253485 | ERS16770978 | - | This study |
| Ba/Apache2 | ocular | Ba | USA | 1960 | ocular | ERR140762 | ERS095032 | - | Andersson et al. 2016 | ERR140762 | ERS095032 | - | Andersson et al. 2016 |
| C/TW3 | ocular | C | Taiwan | 1959 | ocular | ERR558499 | ERS177778 | - | Andersson et al. 2016 | - | - | NC_023057 | Borges et al. 2014 |
| D/SotonD1 | genital | D | UK | 2009 | endocervix | ERR027327 | ERS008761 | - | Harris et al. 2012 | - | - | HE603229 | Harris et al. 2012 |
| D/SotonD6 | genital | D | UK | 2009 | endocervix | ERR027328 | ERS008762 | - | Harris et al. 2012 | - | - | NC_020959 | Harris et al. 2012 |
| D/UW3 | genital | D | USA | 1965 | endocervix | - | - | AE001273 | Stephens Science 1998 | - | - | - | - |
| E/SotonE4 | genital | E | UK | 2009 | endocervix | ERR026551 | ERS013791 | - | Harris et al. 2012 | - | - | NC_020987 | Harris et al. 2012 |
| F/SotonF3 | genital | F | UK | 2009 | endocervix | ERR027330 | ERS008764 | - | Harris et al. 2012 | - | - | HE603234 | Harris et al. 2012 |
| G/SotonG1 | genital | G | UK | 2009 | endocervix | ERR026560 | ERS013800 | - | Harris et al. 2012 | - | - | NC_020961 | Harris et al. 2012 |
| H/R31975 | genital | H | Russia | 2010 | endocervix | ERR111606 | ERS082923 | - | Andersson et al. 2016 | ERR111606 | ERS082923 | - | Andersson et al. 2016 |
| Ia/SotonIa1 | genital | I-Ia | UK | 2009 | endocervix | ERR026555 | ERS013804 | - | Harris et al. 2012 | - | - | NC_020962 | Harris et al. 2012 |
| Ia/SotonIa3 | genital | I-Ia | UK | 2009 | endocervix | ERR026565 | ERS013805 | - | Harris et al. 2012 | - | - | NC_020989 | Harris et al. 2012 |
| J/6276 | genital | J | - | - | endocervix | - | - | ABYD01000001 | Suchland et al. 2008 | - | - | - | - |
| K/SotonK1 | genital | K | UK | 2009 | endocervix | ERR026559 | ERS013799 | - | Harris et al. 2012 | - | - | HE603238 | Harris et al. 2012 |
| L1/115 | LGV | L1 | South Africa | 1986 | urethra | ERR008593 | ERS001411 | - | Harris et al. 2012 | - | - | NC_020951 | Harris et al. 2012 |
| L1/224 | LGV | L1 | USA | 1968 | lymph node | ERR008595 | ERS001396 | - | Harris et al. 2012 | - | - | NC_020952 | Harris et al. 2012 |
| L1/440 | LGV | L1 | South Africa | 1986 | urethra | ERR211058 | ERS161108 | - | Harris et al. 2012 | - | - | NC_010029 | Hatt. 1999 |
| L2/434/Bu | LGV | L2 | USA | 1968 | bubo | - | - | AM884176 | Thomson et al 2008 | - | - | NZ_AM886278 | Thomson et al., 2008 |
| L2b/795 | LGV | L2b | France | 2004 | rectum | ERR008586 | ERS001409 | - | Harris et al. 2012 | - | - | NC_020983 | Harris et al. 2012 |
| L2b/UCH2 | LGV | L2b | UK | - | proctitis | ERR008587 | ERS001405 | - | Harris et al. 2012 | - | - | NC_020956 | Harris et al. 2012 |
| L3/404 | LGV | L3 | USA | 1967 | lymph node | ERR008583 | ERS001416 | - | Harris et al. 2012 | - | - | NC_020957 | Harris et al. 2012 |

**Table S3.** Genes with the highest (Top 1%) number of single nucleotide variant (SNV) accumulation in SvA and SvB sequences.

| CDS | Position (Min) | Position (Max) | SNV |
|---|---|---|---|
| ***Chlamydia trachomatis* SvA** | | | |
| membrane protein CDS | 68,149 | 69,252 | 9 |
| type III secretion system effector protein CDS | 94,989 | 96,671 | 15 |
| ABC transporter substrate-binding protein CDS | 156,524 | 157,804 | 8 |
| DNA ligaseA CDS | 164,295 | 166,286 | 8 |
| hypothetical protein CDS | 166,384 | 170,733 | 19 |
| YitT family protein CDS | 251,627 | 252,523 | 8 |
| inclusion protein (inc) A family CDS | 324,073 | 325,764 | 11 |
| HEAT repeat domain-containing protein CDS | 403,603 | 403,596 | 8 |
| hypothetical protein CDS | 412,199 | 412,735 | 9 |
| translocated actin recruiting phosphoprotein (tarp) CDS | 533,311 | 536,631 | 14 |
| hypothetical protein CDS | 709,788 | 711,743 | 32 |
| SufD family Fe-S cluster assembly protein CDS | 790,296 | 791,483 | 18 |
| hypothetical protein CDS | 847,887 | 849,233 | 8 |
| deubiquitinase CDS | 1,026,820 | 1,028,076 | 15 |
| ***Chlamydia trachomatis* SvB** | | | |
| translation initiation factor IF-2 (infB) CDS | 110,591 | 113,269 | 4 |
| hypothetical protein CDS | 166,347 | 170,696 | 5 |
| inclusion protein (inc) A family CDS | 256,946 | 257,534 | 5 |
| translocated actin recruiting phosphoprotein (tarp) CDS | 533,228 | 536,524 | 4 |
| rpsC CDS | 595,062 | 595,736 | 5 |
| UvrD-helicase domain-containing protein CDS | 689,703 | 691,607 | 4 |
| DNA primase (dnaG) CDS | 931,093 | 932,880 | 4 |
| polymorphic membrane protein (pmp) D CDS | 953,168 | 957,763 | 6 |

**Table S4.** Genes with the highest (Top 1%) number of nonsynonymous to synonymous single nucleotide variant (SNV) among SvA and SvB sequences.

| **CDS** | **Position (Min)** | **Position (Max)** |
|---|---|---|
| ***Chlamydia trachomatis* SvA** | | |
| membrane protein CDS | 68,149 | 69,252 |
| npt1 CDS | 77,379 | 78,959 |
| type III secretion system effector protein CDS | 94,989 | 96,671 |
| YitT family protein CDS | 251,627 | 252,523 |
| hypothetical protein CDS | 274,398 | 275,593 |
| hypothetical protein CDS | 281,737 | 282,087 |
| biotin transporter BioY CDS | 412,795 | 413,385 |
| secD CDS | 521,627 | 525,828 |
| translocated actin recruiting phosphoprotein (tarp) CDS | 533,311 | 536,631 |
| ***Chlamydia trachomatis* SvB** | | |
| infB CDS | 110591 | 113269 |
| hypothetical protein CDS | 166,347 | 170,696 |
| DUF3491 domain-containing protein CDS | 191,189 | 194,041 |
| trpB CDS | 195,097 | 196,275 |
| CTP synthase CDS | 206,611 | 208,203 |
| MFS transporter CDS | 262,582 | 265,368 |
| translocated actin recruiting phosphoprotein (tarp) | 533228 | 536524 |
| rpsC CDS | 595,062 | 595,736 |
| UvrD-helicase domain-containing pr CDS | 689703 | 691607 |
| RNA polymerase factor sigma-54 CDS | 691,611 | 692,921 |
| FAD-dependent thymidylate synthase CDS | 720,998 | 722,587 |
| dnaG CDS | 931093 | 932880 |
| glmM CDS | 959,813 | 961,189 |

**Table S5.** Genes with the highest (Top 1%) number of single nucleotide polymorphism accumulation (SNP) over a short timeframe (4-20 weeks).

***Chlamydia trachomatis*** **SvA/SvB**

| CDS | Position (Min) | Position (Max) | SNP |
|---|---|---|---|
| isoleucyl-tRNA synthetase CDS | 22,026 | 25,136 | 4 |
| class I SAM-dependent methyltransferase CDS | 151,080 | 151,883 | 6 |
| phospholipase D-like domain-containing protein CDS | 181,988 | 183,184 | 3 |
| hypothetical protein CDS | 185,044 | 185,545 | 3 |
| hypothetical protein CDS | 197,792 | 197,962 | 3 |
| small cysteine-rich outer membrane protein CDS | 516,100 | 516, 366 | 3 |
| translocated actin recruiting phosphoprotein (tarp) CDS | 533,228 | 536,524 | 5 |
| deubiquitinase CDS | 1,026,714 | 1,027,970 | 3 |
